# Tissue-autonomous immune response regulates stress signalling during hypertrophy

**DOI:** 10.1101/787226

**Authors:** Robert Krautz, Dilan Khalili, Ulrich Theopold

**Author notes:** Corresponding authors: R. Krautz, U. Theopold. The Bioinformatics Centre, Department of Biology, University of Copenhagen, Ole Maaløes Vej 5, 2200 Copenhagen N, Denmark (Robert Krautz). These authors contributed equally to this work (Robert Krautz, Dilan Khalili).

## Abstract

Postmitotic tissues are incapable of replacing damaged cells through proliferation, but need to rely on buffering mechanisms to prevent tissue disintegration. By constitutively activating the Ras/MAPK-pathway via *Ras^V12^*-overexpression in the postmitotic salivary glands of *Drosophila* larvae, we overrode the glands adaptability to growth signals, induced hypertrophy and stress accumulation. This allowed us to decipher a novel, spatio-temporally regulated interaction between the JNK-stress response and a genuine tissue-autonomous immune response. Central to this interaction is the direct inhibition of JNK-signalling by the antimicrobial peptide Drosomycin, which blocks programmed cell death and prevents recognition of the stressed tissue by the systemic immune response. While this mechanism might allow growing salivary glands to cope with temporary stress, continuous expression of Drosomycin favors survival of unrestricted, hypertrophic *Ras^V12^*-glands. Our findings indicate the necessity for refined therapeutic approaches that fundamentally acknowledge detrimental effects that stimulated immune responses have on tissues coping with damage and stress.

## Introduction

Immune and stress responses have evolved to protect the organism from both exogenous and endogenous stimuli (Eming, 2014; Adamo, 2017; Rankin and Artis, 2018). By sensing deviations from homeostasis and inducing compensatory mechanisms, immune and stress responses keep physiological parameters within tolerable limits (Hoffmann and Parsons, 1991; Vermeulen and Loeschcke, 2007).

Heat shock, radiation, starvation, toxic metabolites, hypoxia or hyperoxia are all well-characterized stressors. They activate stress responses which can ultimately lead to the induction of programmed cell death (Lowe et al., 2004; Loboda et al., 2016). In *Drosophila*, the Keap1-Nrf2-, JNK- and p38-signaling pathways are crucial for mounting these anti-stress-responses (Stronach and Perrimon, 1999; Fuse and Kobayashi, 2017). Immune responses on the other hand, like the Drosophila Toll and Imd pathways, are typically activated by molecular structures exposed on the surfaces of pathogens (Medzhitov and Janeway, 2002; Kurata, 2004). These pathways coordinate the humoral and cellular immune system to eliminate intruding pathogens (Lemaitre and Hoffmann, 2007; Buchon et al., 2014).

Humoral immune responses in *Drosophila* are characterized by the production and secretion of large sets of effector molecules, most notably antimicrobial peptides (AMPs) like Drosomycin (Drs) (Imler and Bulet, 2005). AMPs not only target extrinsic threats, but also react to intrinsic stimuli such as tumorigenic transformation with the possibility to induce apoptosis (Araki et al., 2018; Parvy et al., 2019). However, it remains unknown whether AMPs also have functions beyond promoting apoptosis when sensing and reacting to accumulating stress such as during wound healing and tumor formation.

Apart from their described individual roles, immune and stress pathways are proposed to be either induced successively or concomitantly dependent on the level of deviation from homeostasis (Chovatiya and Medzhitov, 2014; Ammeux et al., 2016). However, detailed characterisation of wound healing and tumor models in *Drosophila* revealed a more complex picture (Park et al., 2004; Buchon et al., 2009; Meyer et al., 2014; Liu et al., 2015; Wu et al., 2015). Accordingly, immune and stress responses often neither occur separately, nor do they follow a simple linear cascade, but rather regulate each other via context-dependent mutual crosstalk (Liu et al., 2015; Liu et al., 2015; Fogarty et al., 2016; Perez et al., 2017). One recurring motif throughout most of these models is the central role of the stress-responsive JNK-pathway and its frequent interaction with the Toll and Imd immune pathways (Rämet et al., 2002; Galko and Krasnow, 2004; Park et al., 2004; Uhlirova et al., 2005; Igaki et al., 2006; Andersen et al., 2015; Enomoto et al., 2015). However, while JNK-signalling can function either in a tumor-promoting, anti-apoptotic or in a tumor-suppressive, pro-apoptotic manner depending on the context, Toll- and Imd-signalling have only been shown to display a tumor-suppressing, pro-apoptotic role in *Drosophila* (Uhlirova et al., 2005; Igaki et al., 2006; Uhlirova and Bohmann, 2006; Vidal, 2010; Cordero et al., 2010; Enomoto et al., 2015).

These tumor-suppressive, pro-apoptotic functions of immune responses have been well characterized and attributed to the secretion of humoral factors or the recruitment of immune cells through the systemic immune system (Babcock et al., 2008; Pastor-Pareja et al., 2008; Hauling et al., 2014; Parisi et al., 2014; Fogarty et al., 2016; Perez et al., 2017). In addition, during clonal cell competition in imaginal discs, Toll- and Imd-signalling were implicated in the elimination of less fit cell clones by inducing apoptosis (Meyer et al., 2014). Importantly, the selective growth disadvantage of these less fit cells is thought to be a response to systemic infection (Germani et al., 2018) and it remains an open question whether genuine tissue-autonomous immune responses can contribute to adaptation of growth during wound healing and tumor formation.

The larval salivary gland (SG) of *Drosophila* is a powerful system to study adaptive growth control, since growth is not completely predetermined, but modulated by the nutritional status (Smith and Orr-Weaver, 1991; Britton and Edgar, 1998). In contrast to mitotically active tissues, growth in post-mitotic tissues like the larval SG is based on endoreplication and hypertrophy rather than on cell division (Edgar et al., 2014; Orr-Weaver, 2015). Modifying the underlying, tight growth regulation, for instance by continuous growth signalling via constitutively activated Ras/MAPK-signalling can easily lead to the accumulation of oxidative stress and DNA damage (Mason et al., 2004; Bartkova et al., 2005; Bartkova et al., 2006; Di Micco et al., 2006; Shim et al., 2013; Shim, 2015). However, the parameters defining the natural limit of growth adaptation and the buffering mechanisms in place to cope with prolonged or continuous stress remain poorly understood.

Here, we uncover a genuine tissue-autonomous immune response which directly regulates hypertrophic growth and adaptation to accumulating stress in larval SGs. By overexpressing a dominant-active form of the small GTPase Ras, *Ras^V12^*, we induced hypertrophic growth. This activates a tissue-autonomous immune response which allows the hypertrophic gland to cope with the resulting stress through spatio-temporally regulated inhibition of the JNK-mediated stress response. We present evidence that tissue-autonomous expression of the AMP Drosomycin (Drs) is at the core of this inhibition: Drs directly interferes with JNK-signalling and inhibits JNK-dependent programmed cell death. This prevents recognition of the stressed tissue by the systemic immune response and allows survival and unrestricted growth of hypertrophic salivary glands.

## Results

### Local immune reaction accompanies Ras^V12^-dependent hypertrophy

In order to identify buffering mechanisms that compensate for continuous stress, we made use of our previously published hypertrophy model in the SGs of *Drosophila* larvae (Hauling et al., 2014). We expressed a dominant-active form of Ras, *Ras^V12^*, uniformly across the entire secretory epithelium of SGs throughout larval development by using the *Bx^MS1096^* enhancer trap (FigS1A for 96/120 h after egg deposition, AED). To further enhance *Ras^V12^*-dependent hypertrophy, we combined *Ras^V12^*-expression with RNAi-mediated knockdown of the cell polarity gene *lethal (2) giant larvae* (lgl; FigS1;(Jacob et al., 1987; Strand et al., 1994)). Their individual and cooperative role in tumor formation in mitotic tissues has been well characterized (Bilder et al., 2000; Pagliarini and Xu, 2003; Herranz et al., 2016). Cell- and tissue-morphology was assessed using Phalloidin-rhodamine staining (Fig1A/S1A) and nuclear morphology by DAPI (Fig1B-C/S1B-D).

**Figure 1.**
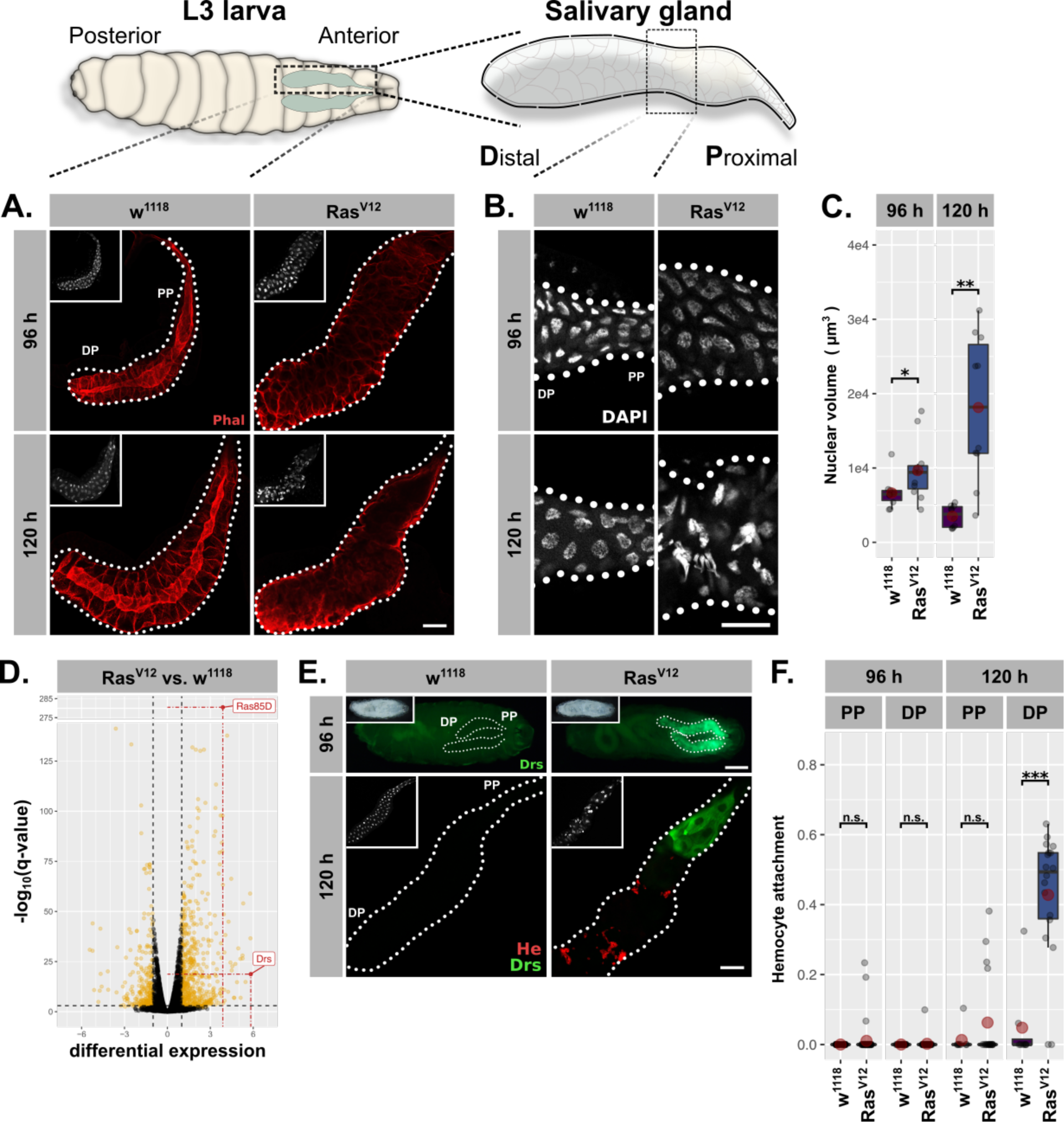
*Ras^V12^*-induced hypertrophy induces local and systemic immune responses. (A.) *Ras^V12^*-glands and controls stained with Phalloidin (red) to monitor tissue integrity at 96 h and 120 h AED. (B.) Nuclei stained with DAPI (white) to visualize nuclear volume and disintegration at 120 h AED in *Ras^V12^*-glands. (C.) Nuclear volume as quantified by z-stacks of DAPI-stained SGs at 96 h and 120 h AED and averaged per gland. (D.) Comparative transcriptome analysis of *Ras^V12^*-vs.-*w^1118^*-glands. Differential expression quantified as beta statistic with q-values by wald test. Significantly differentially expressed genes (*log2(beta)≥1*; *q-value≤0.05)* highlighted in yellow. (E.) Upper: Whole larvae with DrsGFP reporter (green) expressing *Ras^V12^* in glands or controls at 96 h AED. Lower: *Ras^V12^*- and control-glands with DrsGFP reporter (green) stained for hemocytes (anti-Hemese, red). (F.) Hemocyte attachment measured as *ln(Hemese-area)/ln(SG-area)* and separated by time and gland part. Insets: (A./E. Lower) DAPI, (E. Upper) brightfield. Scalebars: (A./B./E. Lower) 100 µm, (E. Upper) 500 µm. Boxplots in (C./F.): lower/upper hinges indicate 1^st^/3^rd^ quartiles, whisker lengths equal 1.5*IQR, red circle and bar represent mean and median. Significance evaluated by Student’s t-tests (*** *p<0.001*, ** *p<0.01*, * *p<0.05, n.s. p≥0.05)*.

At 96 h AED *Ras^V12^*-expressing SG cells retained most of their normal morphology compared to *w^1118^*-control glands. However, their integrity and polarity were severely disrupted at 120 h AED (Fig1A/S1A). Nuclei of *Ras^V12^*-glands showed a continuous increase in volume at 96 h and 120 h AED (1.33 fold compared to *w^1118^* controls at 96 h AED; 5.66 fold at 120 h AED; Fig1C) and signs of nuclear disintegration at 120 h AED implying the induction of programmed cell death (PCD; Fig1B). Both loss of cell integrity and nuclear disintegration coincided temporarily (Fig1A-B), and were exacerbated upon coexpression of *lgl^RNAi^* (FigS1A-D). These results confirm our previous findings and demonstrate that continuous growth signalling in larval salivary glands leads to increased organ size accompanied with additional endocycles at 96 h AED, both hallmarks of compensatory hypertrophy (Hauling et al., 2014: Tamori and Deng, 2014). Furthermore, the additional, *Ras^V12^*-induced endoreplications without obvious effect on tissue integrity imply an adaptability to excess growth signalling, whereas the subsequent collapse of nuclear integrity and cellular polarity at 120 h AED delineate its limitations.

To characterize the mechanisms involved in the early phase of SG growth adaptation at 96 h AED, we performed total RNA sequencing of complete *Ras^V12^*-expressing and *w^1118^*-control SGs dissected at 96 h AED prior to cellular and nuclear disintegration. The most significantly upregulated gene in *Ras^V12^*-glands compared to their *w^1118^*-counterpart was Ras85D itself (q-value=6.51×10^-282^), which validates the experimental set-up (Fig1D). The most differentially expressed gene was the AMP Drs, which indicates the activation of a local immune response in *Ras^V12^*-glands. To evaluate this further, we employed a GFP reporter for Drs and observed strong induction in *Ras^V12^*-glands, but not in any other larval tissue (Fig1E/S1E; Ferrandon et al., 1998a). At 96 h AED the entire secretory epithelium of the SG expressed Drs with a strong tendency for increased induction in the proximal part (PP) closest to the duct (Fig1E). At 120 h AED Drs was almost exclusively expressed in the PP (Fig1E/S1E). In order to assess whether immune cells were recruited as part of a parallel systemic response, we stained glands with an antibody against the pan-hemocyte-marker Hemese. While *Ras^V12^*-glands were completely devoid of hemocytes at 96 h AED, at 120 h AED they were recruited to the gland surface. However, hemocyte attachment was restricted to the distal, non-Drs expressing part (DP), rendering Drs expression and hemocyte attachment across the gland epithelium mutually exclusive (Fig1E-F). Coexpression of *lgl^RNAi^* elevated the level of recruited hemocytes at 120 h AED and pre-empted this recruitment to the DP already at 96 h AED (FigS1E-F).

We next investigated whether the change in nuclear volume as a marker for growth adaptation follows a similar proximal-distal-divide as Drs-expression and hemocyte attachment (FigS1C). Nuclei in the DP of the SG at 96 h AED showed a moderate volume increase upon *Ras^V12^*-expression compared to distal *w^1118^*-control nuclei. However, after 120 h AED distal nuclei had undergone a drastic increase in nuclear volume (6.28 fold compared to distal *w^1118^*-control nuclei) while nuclei in the proximal part of the SG displayed only a moderate increase in size compared to *w^1118^*-control nuclei, that did not increase over time. This indicates that nuclei in the DP of *Ras^V12^*-glands undergo more rounds of endoreplication than their proximal counterparts coinciding with the decline of Drs-expression and an increase in hemocyte attachment in this part.

### Dorsal-dependent Drs expression is part of a genuine tissue-autonomous immune response

As barrier epithelia, the lumen of the SG forms part of a continuum with the exterior, exposing them to extrinsic stimuli including nutritional cues and pathogens (Andrew et al., 2000). Since systemic infection can modulate tissue growth, we sought to clarify whether the observed local immune response in the gland epithelium fulfills the criteria of a genuine tissue-autonomous immune response or was rather embedded in a wider systemic immune response (Germani et al., 2018). Therefore, we eradicated any influence by putative systemic infections, food-derived signals and pathogens or bacterial contamination by raising larvae with *Ras^V12^*-glands under axenic conditions, on minimal medium or by bleaching embryos (FigS2A’-A’’’; Smith and Orr-Weaver, 1991; Britton and Edgar, 1998; Ryu et al., 2008; Piper et al., 2014). None of these changes diminished the Drs expression, strongly indicating that Drs is indeed induced in a *bona fide* tissue-autonomous manner as a response to *Ras^V12^*-dependent hypertrophic growth (FigS2A’-A’’’; Colombani et al., 2005; Mirth et al., 2014).

We further sought to identify the upstream factors controlling Drs expression. The homeobox-like transcription factor Caudal is necessary for Drs expression in the female reproductive organs and the adult SG (Ferrandon et al., 1998a; Han et al., 2004; Ryu et al., 2004). However, RNAi-lines directed against Caudal or its canonical interaction partner, Drifter/vvl, did not reduce Drs-expression in *Ras^V12^*-glands indicating that the regulation of Drs expression as part of the tissue-autonomous immune response is clearly different from its counterpart in the adult SG (FigS2B; Junell et al., 2010).

In order to evaluate whether either Toll- or - as is the case for local infections - Imd-signalling plays a role in Drs expression, we used the reproducible fluorescence pattern of the Drs-GFP reporter at 96 h AED to assay RNAi-lines directed against canonical components of both pathways in *Ras^V12^*-glands (Fig2A; see ‘Materials and methods’ for scored phenotypes; Ferrandon et al., 1998a; Tzou et al., 2000; Takehana et al., 2004; Wagner et al., 2009). Of the 11 RNAi-lines tested, only one targeting the NFκB-transcription factor Dorsal significantly reduced the fluorescence signal of the Drs-reporter (Fig2A). However, Drs expression is completely independent of the upstream modules of the two classical Drosophila immune pathways, Toll and Imd.

**Figure 2.**
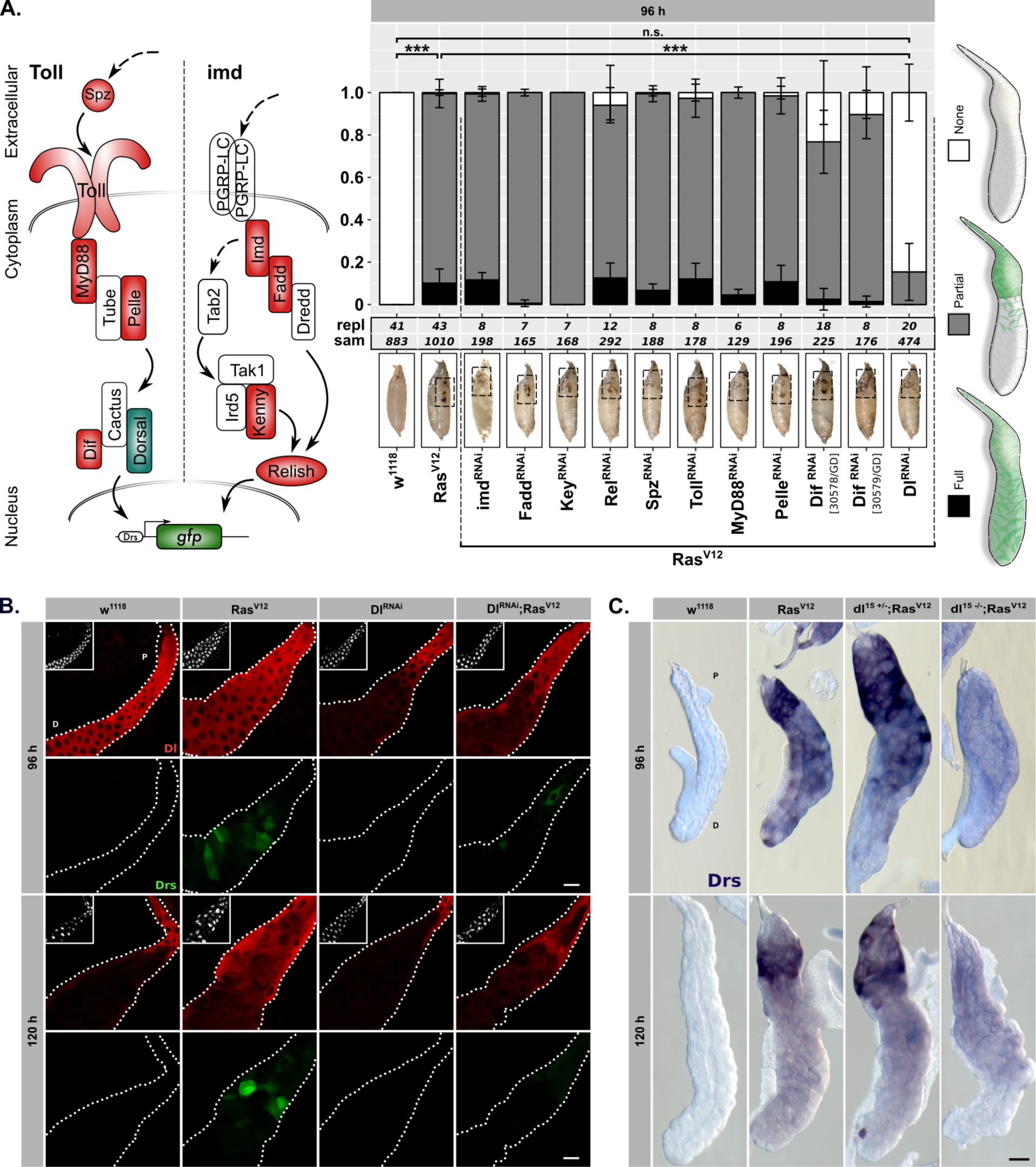
Drs expression is part of a genuine tissue-autonomous immune response. (A.) Semi quantitative DrsGFP reporter assay to identify upstream effectors of Drs expression by RNAi in *Ras^V12^*-glands. Schematic representation (left) of the Toll-/Imd-pathways showing components with (green) and without an effect (red). The 3 distinguished phenotypes (right) were scored per replicate, means and standard deviations plotted (middle) and p-values for “None”-phenotype calculated via Dunn’s test based on Kruskal-Wallis rank sum test (*** *p<0.001*,*n.s. p≥0.05)*. Monitoring melanisation in pupae (insets) confirmed *Ras^V12^*-expression to avoid false positives (Hauling et al., 2014). (B.) Dorsal staining (red) and DrsGFP signal (green) in *Ras^V12^*-glands with and without Dorsal knock-down. (C.) *In-situ* hybridization for endogenous Drs in *Ras^V12^*-glands hetero- or homozygous mutant for Dorsal (dl^15^). Insets: (B.) DAPI. Scalebars: (B./C.) 100 µm.

At 96 h AED Dorsal was present in the entire secretory epithelium of both *Ras^V12^*- and *w^1118^*-control glands. In contrast, at 120 h AED its expression was confined solely to the PP (Fig2B). This overlapped with the Drs-mRNA expression as determined by in situ hybridization (ISH; Fig2SC). Furthermore, both SG-specific knock-down or whole organism homozygous knock-out (dl^15^) of Dorsal completely abolished Drs-expression in the DP of the gland at 96 h AED and drastically reduced it in the proximal part at both time points in line with our results from the reporter assay (Fig2A-B/S2D).

Together, our results indicate that the spatio-temporal dynamics of Drs expression are a consequence of the decrease in endogenous Dorsal expression, independent of canonical Toll- and Imd-signalling. While regulation of Dorsal expression is *Ras^V12^*-independent, Drs is only expressed in the presence of Dorsal during *Ras^V12^*-induced hypertrophy (FigS2E).

### Hypertrophy in SGs induces parallel immune and stress responses

Between 96 h and 120 h AED, the decrease in Drs-expression (Fig1E; Fig2B-C) is correlated with deterioration of tissue integrity (Fig1A-B) in the DP of hypertrophic *Ras^V12^*-glands. Thus, we hypothesized that the tissue-autonomous immune response and Drs-expression aid in preventing the collapse of nuclear as well as cellular integrity until 96 h AED in the DP and until 120h AED in the PP.

To identify possible targets and effector mechanisms of the immune response that buffer the detrimental effects of hypertrophic growth, we analysed the transcriptome data acquired for the *w^1118^*-control and *Ras^V12^*-glands at 96 h AED in further detail. In order to distinguish whether the differences between *Ras^V12^*- and *lgl^RNAi^;Ras^V12^*-glands (FigS1B-F) are of quantitative or qualitative nature and to characterize the PP with its persistent Dorsal and Drs expression in depth, we also profiled transcriptomes of entire *lgl^RNAi^;Ras^V12^*-glands and solely the proximal part of *lgl^RNAi^;Ras^V12^*-glands at 96 h AED (FigS3A; see ‘Materials and methods’).

The sets of significant and differentially upregulated genes (i.e., *log2(beta)≥1*; *q-value≤0.05*) for all three conditions (i.e., *Ras^V12^* / *lgl^RNAi^;Ras^V12^* / *lgl^RNAi^;Ras^V12^*-PP each normalized to *w^1118^*-controls) were intersected to determine common and specific genesets (Fig3A). Notably, the two biggest genesets are differentially upregulated genes shared between all three conditions and genes specifically upregulated in the PP (Fig3A). Furthermore, while all 6 *Ras^V12^*- and *lgl^RNAi^;Ras^V12^*-replicates are in close proximity along the first two principle components in the PCA, the sets of replicates for *w^1118^* and especially the PP are very distant from the rest emphasizing the distinctiveness of the PP compared to the rest of the gland (Fig3B). By screening for enriched gene ontology (GO) terms amongst the significantly upregulated genes in the PP of *lgl^RNAi^;Ras^V12^*-glands in comparison to the entire *w^1118^*-glands, we identified genes belonging to the GO-term ‘immune response’ (i.e., GO:0006955) as significantly enriched (Fig3C; p-value=1.45×10^-4^). Detailed examination of the fold-enrichment of these genes across all three experimental groups (i.e., *Ras^V12^* / *lgl^RNAi^;Ras^V12^* / *lgl^RNAi^;Ras^V12^- PP*) showed a preferential expression in the PP, too (17 of 30 genes highest expressed in PP; blue arrowheads in Fig3C). In addition, 5 of the top 20 upregulated genes in the PP belong to this GO-term as well. Thus, the Drs-expression we observed using reporter lines serves as a proxy for a more complex, tissue-autonomous immune response in hypertrophic glands, especially in the PP. Nonetheless, Drs itself remains one of the top significantly, differentially upregulated genes across all three conditions (Fig1D; FigS3C; not shown for *lgl^RNAi^;Ras^V12^*). In fact, Drs-expression in the PP compared to the two conditions for entire glands is even further increased, confirming the strong tendency for proximal over distal Drs-GFP reporter activation (Fig1E; Fig2B).

**Figure 3.**
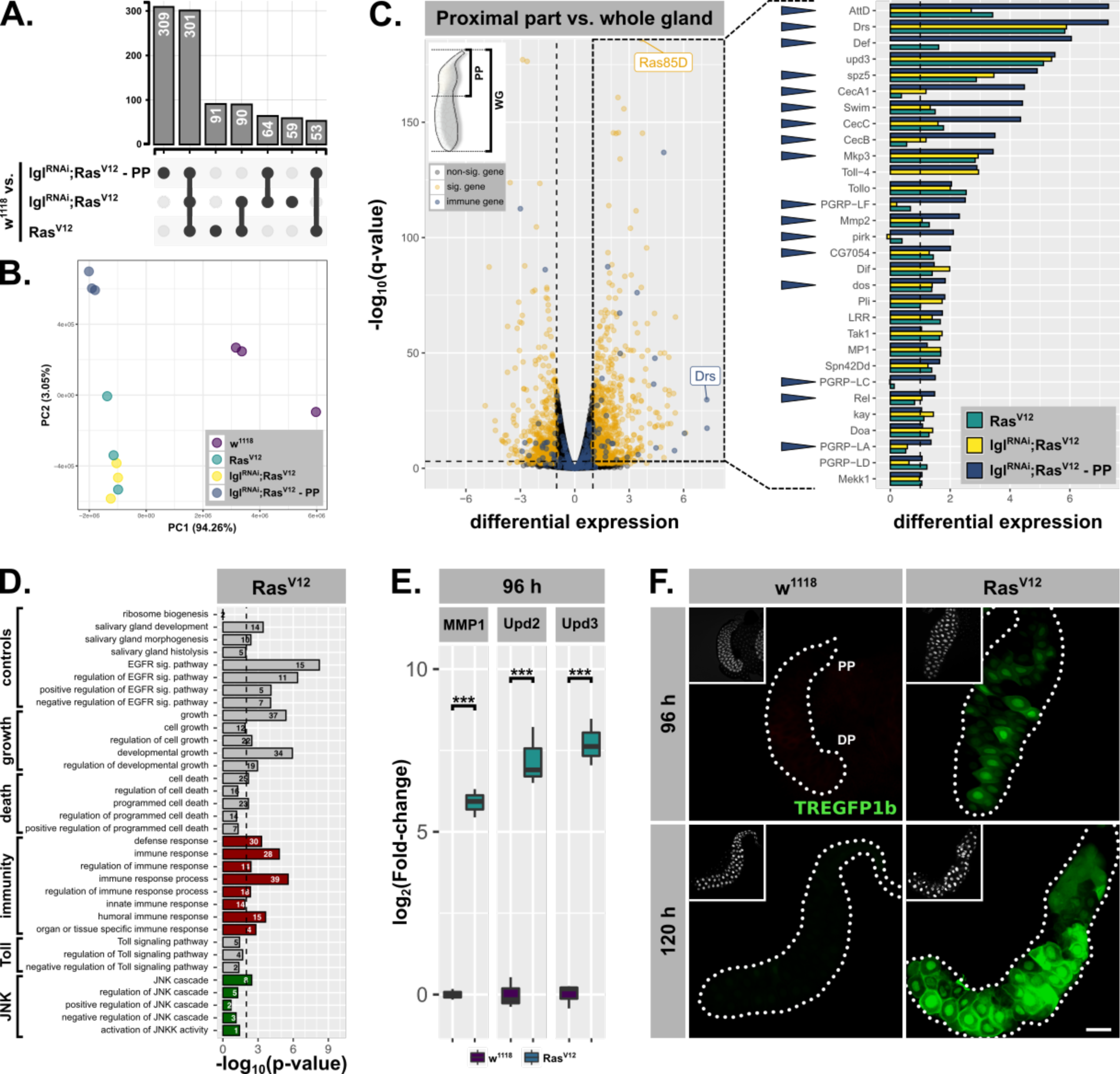
Hypertrophic *Ras^V12^*-glands induce parallel immune and stress responses. (A.) Common and specific genesets significantly upregulated (*log2(beta)≥1*; *q-value≤0.05*) in either *Ras^V12^*, *lgl^RNAi^;Ras^V12^* or *lgl^RNAi^;Ras^V12^*-PP compared to *w^1118^*-glands. (B.) PCA including all transcriptome replicates of all sequenced genotypes. (C.) Left: Comparative transcriptome analysis between PP of *lgl^RNAi^;Ras^V12^*- and entire *w^1118^*-glands. Significantly differentially expressed genes (*log2(beta)≥1*; *q-value≤0.05)* and genes belonging to GO-term ‘immune response’ (GO:0006955) highlighted in yellow and blue. Right: Gene expression in *Ras^V12^*, *lgl^RNAi^;Ras^V12^* or *lgl^RNAi^;Ras^V12^*- PP compared to *w^1118^*-glands for immune genes significantly upregulated in the PP. Blue arrows indicate strongest expression in the PP for the indicated genes between all three groups. (D.) GO term enrichment among significantly upregulated genes in *Ras^V12^*-glands including terms related to activation of JNK (green) and immune responses (red). Numbers in bars indicate amount of upregulated genes belonging to associated GO term. (E.) qPCR results for canonical JNK target genes (*log2*-transformed, fold-change over *Rpl32*) at 96 h AED. Lower/upper hinges of boxplots indicate 1^st^/3^rd^ quartiles, whisker lengths equal 1.5*IQR and bar represents median. Significance evaluated by Student’s t-tests (*** *p<0.001).* (F.) TREGFP1b reporter (green) signal in *Ras^V12^*- and control-glands at 96 h and 120 h AED. Scalbar: 100 µm.

A similar GO-term analysis amongst significantly, upregulated genes in entire *Ras^V12^*- or *lgl^RNAi^;Ras^V12^*-glands revealed the enrichment of genes associated with GO-terms related to ‘growth’, ‘salivary gland development’ and ‘EGFR signalling’ consistent with the studied tissue and the induced *Ras^V12^*-overexpression. In addition, the lack of unique, significantly enriched GO terms and thus distinct gene expression signatures for either *Ras^V12^*- or *lgl^RNAi^;Ras^V12^*-glands at 96 h AED excludes qualitative differences between these two genotypes as an explanation for their phenotypic differences (FigS1).

Importantly, we detected signatures of an activated JNK-cascade as well as cell death in *Ras^V12^*- and *lgl^RNAi^;Ras^V12^*-transcriptomes indicating the presence of an activated stress response and confirming the stimulation of PCD as implied by nuclear disintegration at 120 h AED (Fig1B;Fig3D/S3B GO:0006955; GO:0008219). To validate these signatures further, we performed qPCR for canonical JNK-targets at 96 h AED and found that the expression of all tested genes was significantly increased compared to *w^1118^*-control glands (Fig3E/S3C). We used the TRE-GFP1b-reporter construct, which recapitulates JNK-activation by expressing GFP under the control of binding sites for JNK-specific AP-1 transcription factors (Fig3E; Chatterjee and Bohmann, 2012). This not only confirmed JNK-signalling in *Ras^V12^*-glands, but also uncovered its prevalence in the DP of these glands at 96 h and 120 h AED consistent with the transcriptome data (Fig3E/S3C). Moreover, the elevated fluorescence signal in the DP at 120 h AED compared to 96 h AED implies an increase in activation over time correlating with the collapse of nuclear and cellular integrity during this period (Fig3E/S3C).

In summary, the transcriptome analysis confirms our findings that *Ras^V12^*-overexpression induces a strong tissue-autonomous immune response in the PP of the SG, beyond sole Drs expression. In contrast, the DP shows a striking increase in JNK-signalling which correlates with decreasing Drs expression and cellular and nuclear disintegration at 120 h AED, consistent with the described role of JNK target genes in PCD.

### Drs overexpression and JNK inhibition prevent Ras^V12^-induced tissue disintegration

The increase in JNK-signalling in the DP of the SG as revealed by our transcriptome analysis coincided in space and time with the downregulation of Drs suggesting an interaction between the tissue-autonomous immune response and the stress response.

To test this assumption, we first overexpressed either Drs or a dominant negative form of the *Drosophila* Jun kinase basket individually with *Ras^V12^* throughout the entire secretory epithelium of the SG (Fig4A). Either Drs overexpression or JNK inhibition had a profound effect on the gland size, which was significantly increased at 120 h AED, compared to *Ras^V12^* only (Fig4B/S4A).

**Figure 4.**
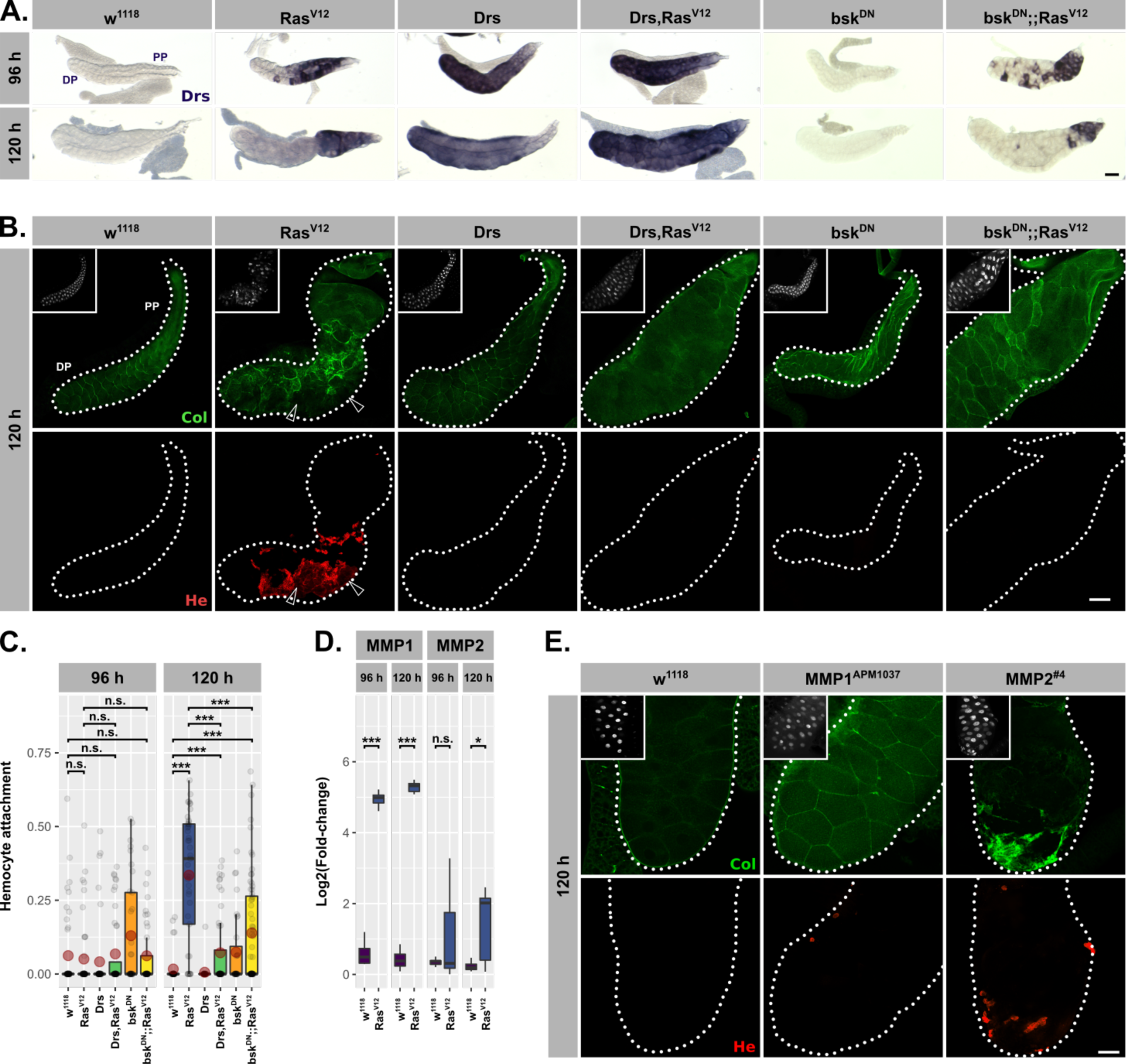
Drs overexpression and JNK inhibition individually prevent tissue disintegration. (A.) Drs-specific *in-situ* hybridization identifies endogenous (*Ras^V12^* / *bsk^DN^;;Ras^V12^*) and exogenous (*Drs* / *Drs,Ras^V12^*) Drs expression. (B.) Collagen-GFP trap (*vkg^G00454^*, green) and Hemese staining (red) identify integrity of BM and hemocyte attachment to gland surface. Arrows indicate BM-free areas occupied by hemocytes. (C.) Hemocyte attachment at 96 h and 120 h AED represented as *ln(Hemese-area)/ln(SG-area)*. (D.) *log2*-transformed, *Rpl32*-normalized gene expression values obtained by qPCR at 96 h and 120 h AED. (E.) Visualization of BM by Collagen-GFP trap (green) and Hemese-stained hemocytes (red) attached to the gland surface upon sole MMP1- or MMP2-overexpression. Insets: (B./E.) DAPI. Scalebars: (A./B./E.) 100 µm. Boxplots in (C./D.): lower/upper hinges indicate 1^st^/3^rd^ quartiles, whisker lengths equal 1.5*IQR, red circle and bar represent mean and median. Significance evaluated by Student’s t-tests (*** *p<0.001*, ** *p<0.01*, * *p<0.05, n.s. p≥0.05)*.

Importantly, despite this size increase, glands of both genotypes (i.e., *Drs,Ras^V12^* / *bsk^DN^;;Ras^V12^*) showed no morphological abnormalities and resembled control *w^1118^*-much more than *Ras^V12^*-glands (Fig4A-B). Strikingly, *bsk^DN^;;Ras^V12^*- and *Drs,Ras^V12^*-glands were completely devoid of attached hemocytes (Fig4B-C). Given the basement membrane’s (BM) role in directly regulating organ morphology, the rescue of the gland shape upon coexpression of either Drs or bsk^DN^ with Ras^V12^ pointed towards changes in the integrity of the BM (Ramos-Lewis and Page-McCaw, 2019). In addition, previous reports suggested that hemocytes are only recruited to tissue surfaces upon tissue disintegration and when the integrity of the BM is lost (Kim and Choe, 2014). Hence, to trace the BM we used an endogenous GFP-trap in the *viking* gene, which encodes one subunit of CollagenIV (Fig4B-C). Both in the presence of Drs or by inhibiting basket, the BM remained a continuous sheet surrounding the entire gland, whereas the BM on the surface of *Ras^V12^*-glands was clearly disrupted.

Matrix metalloproteinases (MMPs) are likely candidates for executing the disruption of the BM and known target genes of the JNK-pathway (Hauling et al., 2014; Stevens and Page-McCaw, 2012; Uhlirova and Bohmann, 2006). In order to see whether hypertrophic glands express MMPs, we performed qPCR for MMP1 and MMP2 at 96 h and 120 h AED on *Ras^V12^*- and control *w^1118^*-glands. At 96 h AED, *Ras^V12^*-glands already exhibit increased MMP1 expression (Fig4C-D). However, MMP2 only reaches a significant level of expression at 120 h AED coinciding with the appearance of hemocyte attachment (Fig4D). SG-wide, *Ras^V12^*-independent overexpression of MMP2, but not of MMP1, caused opening of the BM and hemocyte attachment to the surface (Jia et al., 2014). This indicates the necessity for a JNK-dependent expression of MMP2 in the hypertrophic *Ras^V12^*-glands prior to the recruitment of hemocytes to the tissue surface (Fig4E/S4B).

Taken together, overexpression of Drs alone is sufficient to mimic the inhibition of the JNK-pathway in *Ras^V12^*-glands: both lead to excess hypertrophic growth compared to *Ras^V12^*-glands, but simultaneously prevent tissue disintegration and PCD. Prohibiting the disruption of the basal membrane inhibits hemocyte recruitment and thus prevents the systemic immune response from sensing hypertrophic growth. This in turn suggests that the endogenous Drs expression in *Ras^V12^*-glands is seminal for maintaining nuclear and tissue integrity and thus part of the buffer mechanism to adapt to continuous growth signalling.

### Drs inhibits JNK-signalling

The strong correlation between loss of Drs and the increase in JNK-signalling in the DP of *Ras^V12^*-glands between 96 h and 120 h AED indicated an active interaction between Drs and the JNK-pathway, which prompted us to resolve their hierarchy by epistatic analysis.

Coexpression of *bsk^DN^* with *Ras^V12^* did not deplete Drs in the proximal part of 120 h-old glands, since neither the fluorescence signal of the Drs-GFP reporter nor staining for endogenous *Drs*-mRNA via ISH showed any effects, which excludes a direct regulation of Drs by JNK-signalling (Fig4A;FigS5A). qPCR for Drs in *Ras^V12^*- and *bsk^DN^*;*Ras^V12^*-glands confirmed these results further (FigS5B). In contrast, qPCR in hypertrophic *Ras^V12^*-glands showed a significant reduction in expression of JNK target genes upon coexpression of Drs at 96 h and 120 h AED (Fig5B/S5C-D), in line with a decrease in activated basket (Fig5C,E) and TRE-GFP1b signal in *Drs,Ras^V12^*-compared to *Ras^V12^*-glands (Fig5B,D,F). Thus, the overexpression of the AMP Drs actively and tissue-autonomously inhibits the JNK-dependent stress response in hypertrophic *Ras^V12^*-glands beyond 96 h AED.

**Figure 5.**
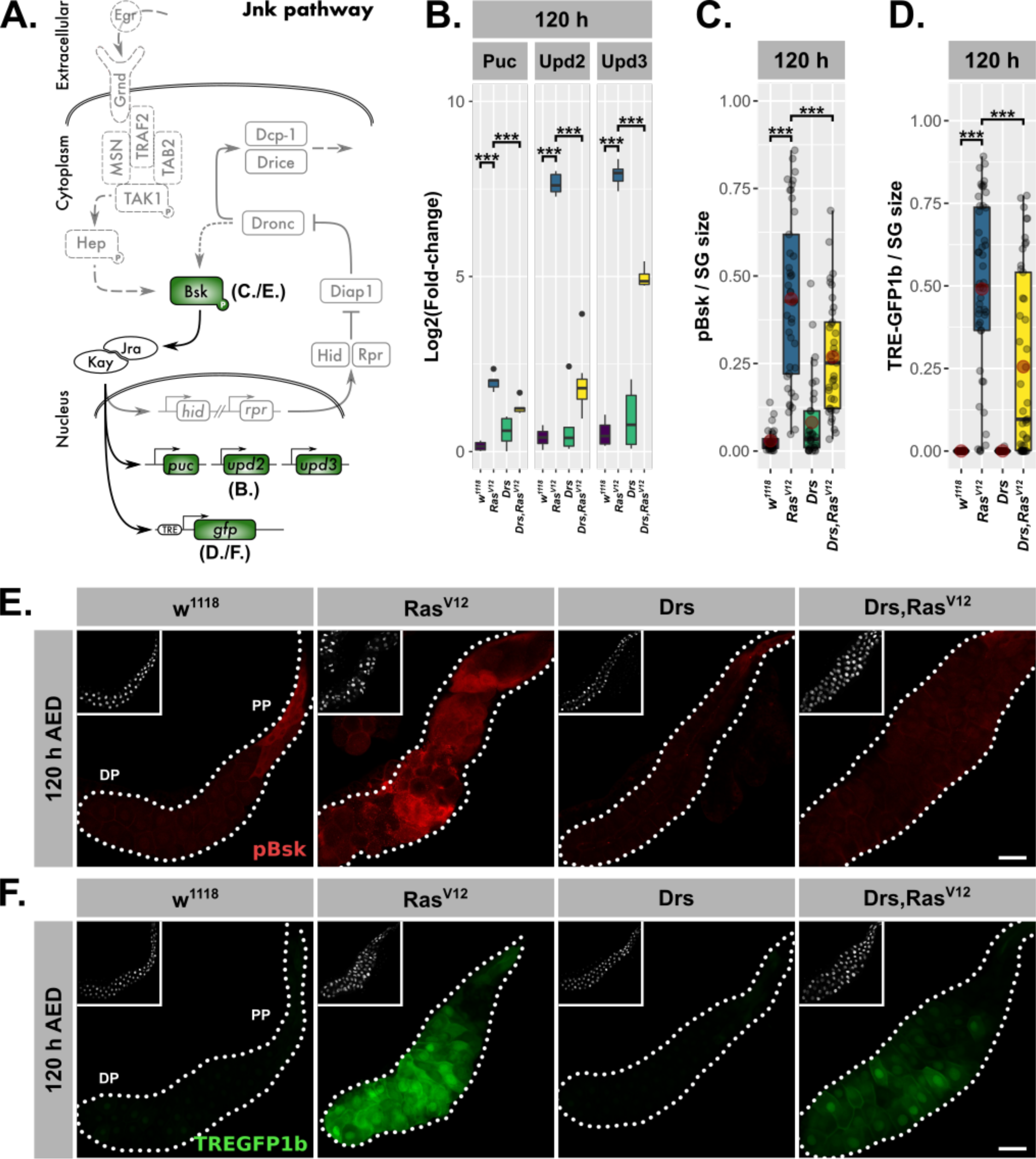
Drs overexpression inhibits JNK-activation. (A.) Schematic representation of the JNK-pathway including read-outs (green) employed to track its activation. (B.) qPCR results for canonical JNK-target genes (*log2*-transformed, fold-change over *Rpl32*) at 120 h AED. (C.) Quantification of activated basket by staining with phosphorylation sensitive antibody per gland normalized for the SG size at 120 h AED. (D.) Quantification of TREGFP1b signal per gland normalized for SG size indicating JNK-dependent transcriptional activation at 120 h AED. (E.) Visualization of phosphorylated basket in *Ras^V12^*-glands with and without Drs coexpression. (F.) Distribution of TREGFP1b reporter signal in *Ras^V12^*-glands in the presence or absence of coexpressed Drs. Insets: (E./F.) DAPI. Scalebars: (E./F.) 100 µm. Boxplots in (B./C./D.): lower/upper hinges indicate 1^st^/3^rd^ quartiles, whisker lengths equal 1.5*IQR, red circle and bar represent mean and median. Significance evaluated by Student’s t-tests (*** *p<0.001*, ** *p<0.01*, * *p<0.05, n.s. p≥0.05)*.

In addition, we used an efficient RNAi-line against Drs, which in combination with Ras^V12^ reduced Drs-expression drastically compared to its expression in *Ras^V12^*-glands (FigS5C-D). All significantly upregulated JNK-target genes in *Ras^V12^*-glands apart from *upd2* were further increased upon knockdown of Drs at 96 h AED (FigS5C). However, this effect ceased at 120 h AED consistent with the canonical loss of Drs expression in the DP of *Ras^V12^*-glands (Fig5D). This result demonstrates that until 96 h AED the endogenously expressed Drs has the same inhibitory effect on the JNK-pathway as the Drs-overexpression has at 120 h AED.

### Drs prevents cell death in Ras^V12^-glands

The capacity to induce apoptosis is a well-established function of the JNK-pathway in *Drosophila* (Uhlirova et al., 2005; Igaki et al., 2006; Uhlirova and Bohmann, 2006; Cordero et al., 2010; Enomoto et al., 2015). Combined with the identification of ‘cell death’ as a significantly enriched term in the GO-analysis, we hypothesized that the observed nuclear disintegration in *Ras^V12^*-glands after 96 h AED was a consequence of the JNK-dependent induction of PCD (Fig1B-C/S1B-D; Fig3D/S3B).

In mitotic tissues, the apoptotic inducer *head involution defective* (hid) is inhibited by Ras/MAPK-signalling (Bergmann et al., 1998; Kurada and White, 1998). However, in *Ras^V12^*-glands hid expression gradually increased from 96 h to 120 h AED coinciding with the increase in nuclear disintegration (Fig6A). Coexpression of Drs with *Ras^V12^* significantly decreased hid expression at both time points, while Drs knock-down in *Ras^V12^*-glands increased hid expression even further already at 96 h AED. Together with the reduction in hid expression upon JNK-inhibition in *bsk^DN^;;Ras^V12^*-glands, Drs emerges as a negative regulator of the apoptotic inducer hid by inhibiting JNK-signalling (FigS5B).

**Figure 6.**
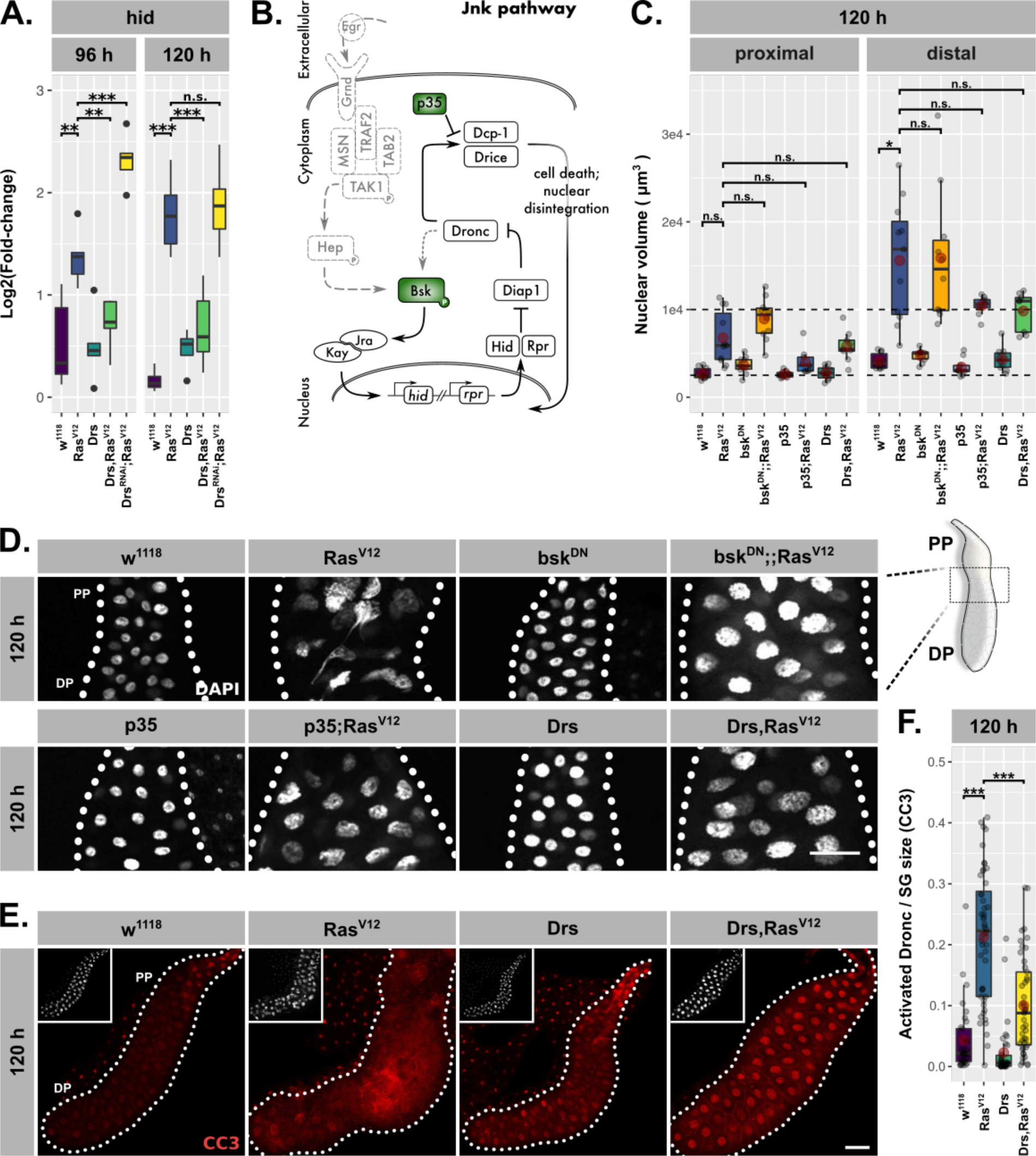
Drs inhibits programmed cell death. (A.) Hid expression as measured by qPCR and plotted as *log2*-transformed values normalized to *Rpl32*-expression in *Ras^V12^*-glands with in- (Drs) or decreased (Drs^RNAi^) Drs expression. (B.) Schematic representation of the JNK-pathway including levels of employed interference (green). (C.) Nuclear volumina derived from z-stacks of DAPI-stained SGs and averaged per gland at 96 h and 120 h AED. (D.) DAPI stained (white) SG nuclei to indicate nuclear size and disintegration. *Ras^V12^*-glands with in- (Drs) or decreased (Drs^RNAi^) Drs expression (E.) stained with anti-CC3-antibody to detect Dronc activation (red) and (F.) corresponding quantifications of detected signal normalized for SG size. Insets in (E.) show DAPI and scalebars in (D./E.) represent 100 µm. Lower/upper hinges of boxplots in (A./C./F.) indicate 1^st^/3^rd^ quartiles, whisker lengths equal 1.5*IQR, red circle and bar represent mean and median. Significance evaluated by Student’s t-tests (*** *p<0.001)*.

To evaluate the induction of PCD in *Ras^V12^*-glands, we either inhibited JNK-signalling at the level of basket activation or coexpressed the caspase-inhibitor p35 with *Ras^V12^* and examined nuclear volume and integrity (Fig6B-D). p35 inhibits Drice and thus blocks PCD at the level of effector caspase activation (Hawkins et al., 2000; Hay et al., 1994; Meier et al., 2000). While both interventions successfully blocked nuclear disintegration, the additional rounds of endoreplication as observed in *Ras^V12^*-glands were not suppressed (Fig6C-D). Crucially, Drs-coexpression in *Ras^V12^*-glands phenocopies the inhibition of JNK-signalling and effector caspases both in terms of restoring nuclear integrity as well as the persistence of excess endoreplications.

Last, to validate the inhibition of PCD by Drs, we monitored Dronc activity using the cleaved caspase 3 antibody (CC3). In fact, the strongest Dronc activity occurred in cells that also displayed heavy disintegration of nuclei, confirming the relation between caspase activation and nuclear disintegration as part of PCD (FigS6A-B). However, in *Drs,Ras^V12^*-glands Dronc was significantly less activated compared to *Ras^V12^*-glands (Fig6E-F; Fan and Bergmann, 2010).

Taken together, we show that JNK-dependent induction of PCD is a consequence of the collapse in adaptation to *Ras^V12^*-induced hypertrophy. Importantly, overexpression of Drs in *Ras^V12^*-glands prevents PCD by blocking full JNK-activation. This is in contrast to recent observations where AMPs, including Drs, act pro-apoptically aiding in the elimination of tumor cells and limiting tumor size (Araki et al., 2018; Parvy et al., 2019).

### Drs inhibits the JNK-feedback loop

Various *Drosophila* models for tissue transformation have shed light on the functional separation of the JNK-pathway into an upstream kinase cascade leading to basket-activation and a downstream feedback-loop converging again on basket activation (Fig7A; Shlevkov and Morata, 2012; Fogarty et al., 2016; Muzzopappa et al., 2017; Perez et al., 2017).

**Figure 7.**
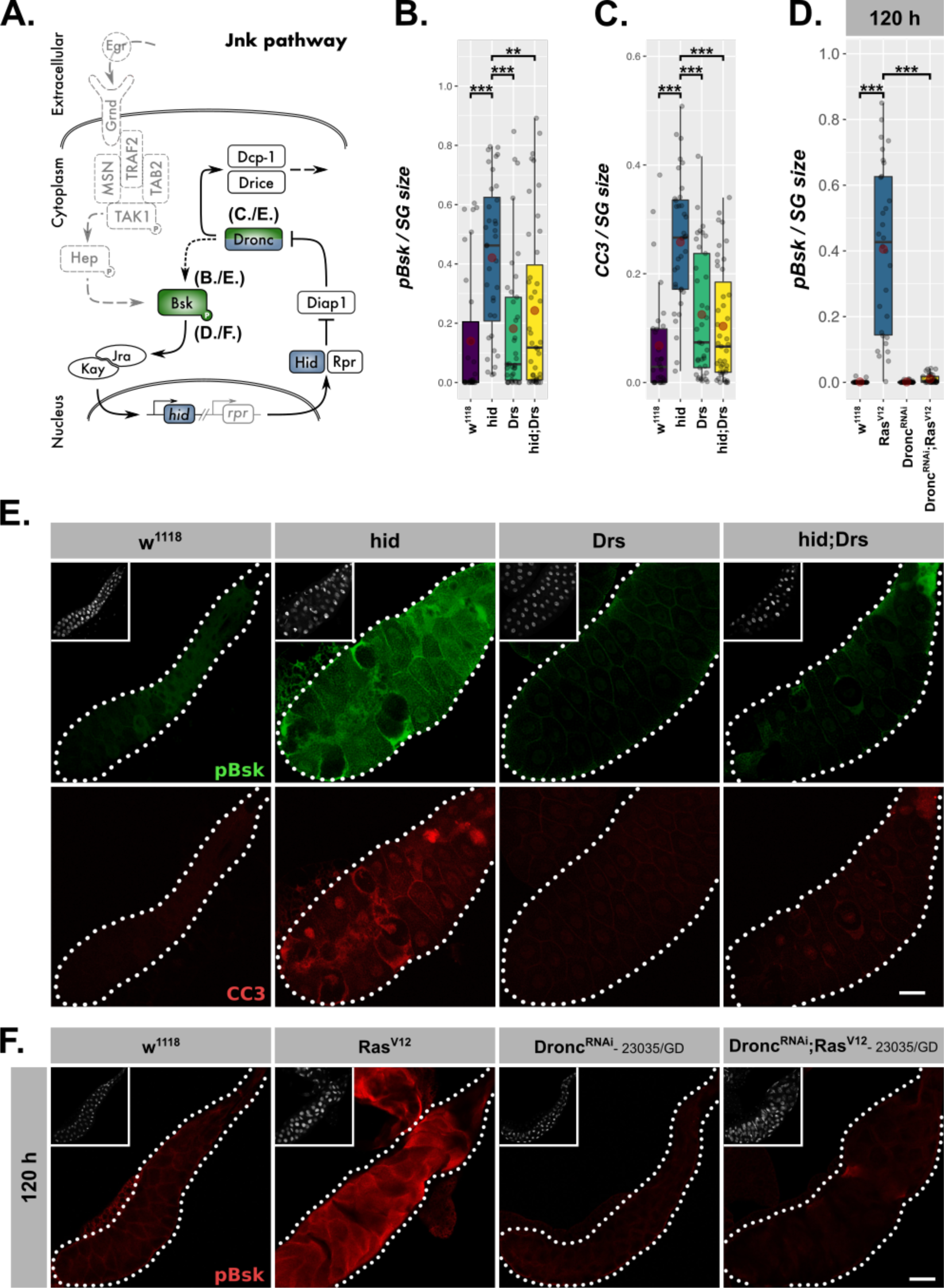
Drs inhibits the JNK-feedback loop. (A.) Schematic representation of the JNK-feedback loop including levels of interference (blue) and employed read-outs of its activity (green). *hid*-overexpressing glands with or without coexpressed Drs were quantified for (B.) activated basket via a phosphorylation sensitive antibody and (C.) activated Dronc via the CC3-antibody. (D.) Activated basket (pBsk) was quantified and normalized to SG size. (E.) *Ras^V12^*-glands with or without Drs-coexpression were stained for activated basket (green) and activated Dronc (red). (F.) Activated basket was detected in *Ras^V12^*-glands with or without Dronc-knock-down. Insets: (E./F.) DAPI. Scalebars: (E./F.) 100 µm. Boxplots in (B./C./D.): lower/upper hinges indicate 1^st^/3^rd^ quartiles, whisker lengths equal 1.5*IQR, red circle and bar represent mean and median. Significance evaluated by Student’s t-tests (*** *p<0.001*, ** *p<0.01*, * *p<0.05, n.s. p≥0.05)*.

Our results revealed a negative regulatory impact of Drs on JNK-signaling in *Ras^V12^*-glands, but did not determine which part of the pathway is targeted by Drs. To answer this question, we uncoupled the feedback loop from the upstream kinase cascade by solely overexpressing hid (Fig7A). Irrespective of the differing models for the signal propagation downstream of Dronc, this initiator caspase remains seminal for establishing the actual feedback with basket (Shlevkov and Morata, 2012; Fogarty et al., 2016). Thus, we stained glands for activated Dronc and basket after a pulse of hid-overexpression during the larval wandering stage (Fig7A). As expected, *hid*-expressing glands showed highly elevated levels of activated Dronc, but also basket, which indicates the presence of feedback activation. Moreover, both phenotypes were reversed upon coexpression of Drs during the hid-expression pulse (Fig7B-C,E). Thus, Drs seems to inhibit JNK-signalling downstream of hid and upstream of Dronc and basket, emphasizing an inhibition of the JNK-feedback loop rather than the initial kinase cascade.

To clarify that Drs operates in a similar fashion during *Ras^V12^*-induced hypertrophic growth, we stained *Ras^V12^*-glands (Fig6E-F) for activated basket in the presence and upon knock-down of Dronc (Fig7D,F/S7A-B). In fact, the absence of a signal for activated basket in *Dronc^RNAi^;Ras^V12^*-compared to strong *Ras^V12^*-glands confirms the presence of a genuine feedback-regulation as part of JNK-signalling in SGs.

## Discussion

### Tissue-autonomous vs. systemic immune response mediated via JNK-signalling

By overexpressing *Ras^V12^* in larval SGs, we made use of and overrode the gland’s ability to adapt to growth signals. Activating constitutive Ras/MAPK-signalling allowed us to identify local immune and stress responses as part of a buffering mechanisms to compensate the accumulation of stress and decipher natural limits of growth adaptation (Hauling et al., 2014). Remarkably, in this context the local immune response inhibits the parallel stress response, an effect that to our knowledge has not been described before. Central to this inhibition is the AMP Drs, which directly impinges on the JNK-pathway and thereby subsequently also on inducing apoptosis, a function completely opposite to previous observations in other tissues (Araki et al., 2018; Parvy et al., 2019). These results also indicate an unprecedented role for an AMP as a signal transducer enabling tissue-autonomous crosstalk between immune and stress pathways. Dependent on the extent of JNK-inhibition by the local, Drs-dependent immune response and thus the integrated decision of both on the state of the tissue’s homeostasis, the gland epithelium attracts hemocytes as part of a wider, systemic immune response. By virtue of inhibiting JNK-signalling, tissue-autonomous and systemic immune response antagonize each other, a balance that decides on continuos hypertrophic growth or its restriction (Fig8). The latter has far reaching implications for therapeutic approaches that need to consider the adverse effects that stimulating immune responses might have on tissue growth after damage and under stress.

**Figure 8.**
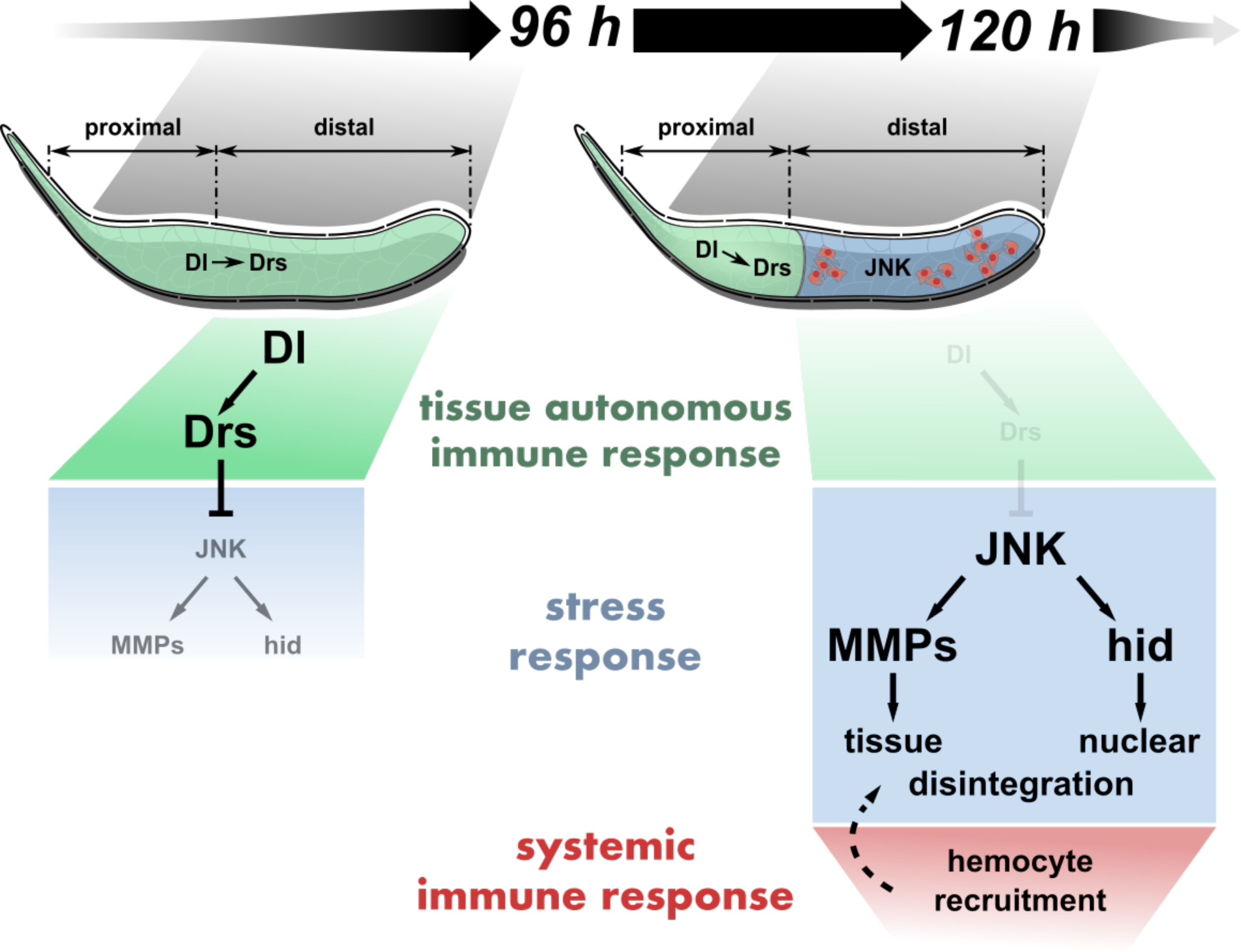
Tissue-autonomous antagonizes systemic immune response via JNK-inhibition in *Ras^V12^*-glands. Dorsal-mediated Drs expression inhibits JNK activation in the entire *Ras^V12^*-gland until 96 h AED. At 120 h AED, Dorsal and thus Drs expression is reduced to the PP, derepressing full JNK-activation in the DP. Consequently, JNK-dependent expression of hid and MMP2 stimulates nuclear and tissue disintegration, which eventually leads to the interference with tissue growth and integrity by the systemic immune response.

### Drs is expressed as part of a genuine tissue-autonomous immune response

The salivary glands of *Drosophila* larvae are an integral part of its gastrointestinal system and the lumen of the mature glands forms a continuum with the exterior. As such the glands are constantly exposed to microbial and pathogenic influences, which predestines them to be a dedicated immunological barrier epithelium. However, raising larvae with *Ras^V12^*-glands under strictly axenic conditions corroborated the authenticity of the immune response as truly tissue-autonomous and thus independent of exogenous or systemically distributed pathogenic stimuli. This is further emphasized by the dependency of Drs expression on the tissue-specific overexpression on *Ras^V12^* and tissue-autonomous manipulations (i.e., Drs or bsk^DN^ overexpression) leading to the inhibition of JNK-activation and hemocyte recruitment.

However, apart from Dorsal neither the homeobox transcription factor Caudal as in the adult glands nor any other of the canonical members belonging to the Toll- and Imd-pathway are involved in Drs expression (Ferrandon et al., 1998b; Hauling et al., 2014; Ryu et al., 2004). Under wild-type conditions, Dorsal is expressed throughout the entire gland epithelium until 96 h AED, but only remains expressed in the PP at 120 h AED (Fig2B). This spatio-temporal expression pattern in turn determines the expression of Drs in *Ras^V12^*-glands essentially separating the gland at 120 h AED into an immunocompetent duct-proximal and a stress-responsive duct-distal compartment (Fig2B,Fig8). Due to the absence of Drs in the DP at 120 h AED, stress-responsive JNK-signalling finally exceeds a critical threshold leading to elevated MMP2- and hid-expression. This stimulates the onset of PCD and opening of the basal membrane as a prerequisite for attachment of hemocytes that are subsequently recruited to the surface of the SGs (FigS2E,Fig8; Hauling et al., 2014). To the contrary, Dorsal continues to induce Drs in the PP of *Ras^V12^*-glands until 120 h AED in line with the complete absence of hemocyte recruitment to this part of the gland (Fig1F/S1B-C).

### Drosomycin impinges on JNK feed-back activation

In depth analysis of wound regeneration and tumor formation has shed light on the intricate architecture of the JNK-pathway and its signal propagation (Santabarbara-Ruiz et al., 2015; Pinal et al., 2018). This has led to the discovery of feedback- or self-sustenance-loops as part of JNK-signalling (Shlevkov and Morata, 2012; Fogarty et al., 2016; Muzzopappa et al., 2017; Perez et al., 2017). Similarly, several lines of evidence validated the existence of a complex quantitatively and qualitatively regulated activation of the JNK-pathway in *Ras^V12^*-glands. In fact, the strong reduction of basket activation upon knock-down of the initiator caspase Dronc in the *Ras^V12^*-background at 120 h AED indicates that a feedforward-loop is also part of the propagation of JNK-activation in hypertrophic glands (Fig7D,F/S7A-B). Consistently, even sole hid overexpression in SGs led to Dronc as well as basket activation emphasizing the presence of this feedback-regulation as part of the JNK-pathway further (Fig7B-C,E).

In general, feedback loops serve as a predestined platform to integrate additional signals via crosstalk with other pathways to dynamically modulate the originally, transmitted signal and thus either amplify or weaken the response (Antebi et al., 2017; Kholodenko, 2006). Here, we describe a novel mode of signal attenuation by the tissue-autonomous immune response that rather than eliminating an exogenous stimulus directly interferes with the signal propagation in the JNK-feedback loop. Fundamentally, this interaction is part of an emerging picture of crosstalk between immune and stress responses involved in organ growth and maintenance in *Drosophila* (Hauling et al., 2014; Liu et al., 2015; Parisi et al., 2014; Wu et al., 2015).

Drs regulates the signal propagation of the intracellular module of the JNK-pathway and throughout our experiments only directly Drs expressing cells prevented *Ras^V12^*-induced cell disintegration and PCD. However, it remains an outstanding question whether Drs functions exclusively cell-autonomously and intracellularly or whether secreted Drs operates in an autocrine manner too. Clonal analysis and rescue experiments will serve this purpose in the future. Further work on the effector mechanism of Drs will also elucidate further details about the components of the JNK-feedback loop in- or directly regulated by Drs. This will also contribute to mapping the manifold modes of immune-stress-crosstalk in *Drosophila* and find general patterns among them beyond a sole dependency on the specific context.

### Drs promotes hypertrophic growth and inhibits PCD

As our *Drs,Ras^V12^*-experiments indicated, under conditions of continuous growth and therefore chronic stress induction, the ability to suppress the JNK-pathway in a Drs-dependent manner supports continuous, hypertrophic growth of *Ras^V12^*-glands and the survival of the tissue. Moreover, a prolongation of Drs-expression beyond its endogenous decrease inhibits the induction of PCD and the recognition of the systemic immune response and thus renders the hypertrophic gland unchallenged.

In fact, while Ras/MAPK-signalling in mitotic tissues crucially suppresses apoptosis by downregulating hid expression, this effect is revoked in hypertrophic *Ras^V12^*-glands (Bergmann et al., 1998; Kurada and White, 1998). While the reason for this difference remains to be elucidated, it is in fact Drs which operates as the inhibitor of apoptotic inducers in hypertrophic glands. This function is central to the suppression of PCD in *Ras^V12^*-glands (Fig6A).

This differs fundamentally from the pro-apoptotic function of AMPs, which was recently described for two tumor models. In both disc (discs large, *dlg,* Parvy et al., 2019) and leukemic tumors (mxc^mbn1^, Araki et al., 2018), AMPs were shown to target tumor cells and limit tumor size by inducing apoptosis. In addition to the differences between the tumorous tissues (i.e., proliferative discs and lymph glands), the fact that Drs is induced locally in SGs and acts tissue-autonomously may explain its different activities.

### Hypertrophic SGs are a remarkable system to discover buffer mechanisms

Being incapable of cell proliferation, damaged postmitotic tissues can’t rely on regenerative cell plasticity like imaginal discs or stem cell-derived tissue regeneration as in the *Drosophila* adult midgut (Ohlstein and Spradling, 2006; Herrera et al., 2013; Herrera and Morata, 2014; Schuster and Smith-Bolton, 2015; Ahmed-de-Prado and Baonza, 2018). Instead, they need to cope with endogenous and exogenous influences via elaborate mechanisms to prevent or buffer detrimental consequences.

Here, we show that continued *Ras^V12^*-expression in the larval salivary gland overrides the dependency on nutritional cues and stimulates excess endoreplications that eventually have damaging effect on the tissue integrity by inducing elevated levels of stress. Uninterrupted induction of endoreplications has its natural limits in every system, even in salivary glands that are already polyploid. Eventually, continuous induction of more endocycles is challenged by nutritional restrictions in synthesizing more DNA, replication stress, spatial limitations in the gland nuclei and continuously more unsynchronized metabolic turnover. Hence, in spite of the anti-apoptotic function *Ras^V12^* conveys in mitotic tissues, unrestricted stimulation of excess endoreplications ultimately leads to cell death. Since this characterizes the final collapse of tissue homeostasis, it also allows to study the extent of buffering capacity conveyed by immune and stress-responsive signalling.

Thus, hypertrophic *Ras^V12^*-glands constitute an outstanding system to study the involvement of tissue-autonomous immune and stress-induced responses to buffer deviation from homeostasis.

### Immune surveillance theory

According to the immune surveillance theory, the immune system has evolved to reduce the risk of somatic cells accumulating cancerous mutations (Burnet, 1970; Burnet, 1957). In order to reduce the danger of cell-transformation, cells express or expose molecules upon recognition of stress or damage during transformation. These markers are sensed by the immune system, which in turn eradicates the potentially harmful cells. (Jung et al., 2012; Vantourout et al., 2014; Schmiedel and Mandelboim, 2018). However, the immune surveillance theory remains controversial, since tumor-associated inflammation was also shown to promote rather than suppress tumor growth (Balkwill and Mantovani, 2001; Mantovani et al., 2008). Our model bridges the gap between these two opposing views, since the effects of the tissue-autonomous and systemic immune responses appear to be antagonistic regarding the regulation of JNK-activation and thus ultimately PCD. In fact, only the integration of the various stress and immune mechanisms in hypertrophic *Ras^V12^*-glands allows a concerted decision to eradicate a putatively dangerous cell via inducing PCD or not. Given the evolutionary conservation from insects to mammals of signalling pathways that govern growth control (Edgar, 2006), it is likely that mechanisms to detect and counteract a loss in regulation of these pathways, such as stress and immune pathways, are similarly conserved between both phyla.

## Material and methods

### Fly husbandry and stocks

All crosses were reared on standard potatomash/molasses medium under tempered conditions (see ‘Staging’) in a 12h dark/12 h light-cycle. Drs-GFP (W.-J. Lee), UAS-dfr^RNAi^ (S.Certel), TRE-GFP1b (D. Bohmann), UAS-Drs (B. Lemaitre), dl^15^ (Y. Engström), UAS-hid (M.Suzanne), UAS-MMP2^#4^, UAS-MMP1^APM1037^ and UAS-MMP1^APM3099^ (A. Page-McCaw) were kind gifts from the indicated donors. The generation of the CollagenIV flytrap line vkg^G00454^ was described in (Morin et al., 2001). Please, see supplementary file 1 for complete list of experimental crosses.

### Staging

Virgins were collected for 3-7 d before setting crosses. Initially, crosses were kept on standard food without antibiotics for 48 h at 25°C. Eggs were collected for 6 h (Immunohistochemistry and qPCR) or for 2 h (RNASeq) at 25°C. When necessary, precollections were performed for 2 h at 25°C prior to the actual collection. Egg collections were incubated for 24 h at 29°C or for hid experiments 48 h at 18°C and batches of 24 hatched 1^st^ instar larvae were afterwards transferred to vials with 3 ml standard food supplemented with Neomycin (0.1 mg·ml^-1^, Sigma-Aldrich, N1876), Vancomycin (0.1 mg·ml^-1^, Sigma-Aldrich, V2002), Metronidazol (0.1 mg·ml^-1^, Sigma-Aldrich, M3761) and Carbenicillin (0.1 mg·ml^-1^, Sigma-Aldrich, C1389). After incubation at 29°C for another 72 h (96 h AED) or 96 h (120 h AED), 3^rd^ instar larvae were prepared for dissection or pictures were taken of whole larvae with a Leica MZ FLIII Fluorescence Stereomicroscope. For hid experiments, transferred larvae were incubated for 197 h at 18°C, shifted to 29°C for 12 h and finally dissected. To exclude microbial contamination and maintain germ-free conditions, BxGal4;;DrsGFP/UAS-Ras^V12^-eggs were dechorionated with a 50% Sodium Hypochlorite solution (Fisher Scientific, 10401841) immediately after collection and transferred to vials with apple-agar supplemented with Nipagin, Propionic acid and the same antibiotics as above. Larvae were then analysed at 24 h and 48 h AED.

### Drs reporter assay

Replicates of *Drosophila* larvae (n=24 larvae·replicate^-1^; N>6 replicates) at 96 h AED were screened for Drs-GFP reporter signals. Three phenotypes were distinguished: (1.) “Full” SG pattern includes GFP signal across the entire PP and GFP^+^-cells with reduced intensity in the DP. (2.) “Partial” SG is GFP^+^ throughout the PP, but less pronounced in DP with reduced signal intensity and fewer GFP^+^-cells. (3.) “None” phenotype lacks signal throughout the entire gland. Distribution of phenotypes were scored per replicate and significance calculated for the “None”-phenotype via Dunn’s test after performing the Kruskal-Wallis rank sum test.

### qPCR

Total RNA of dissected salivary glands was isolated with the RNAqueous-Micro Kit (ThermoFisher Scientific, AM1931) and from whole larvae with the RNAqueous Kit (ThermoFisher Scientific, AM1912) according to the instructors manual. Residual genomic DNA was digested with RNase-free DNaseI (ThermoFisher Scientific, EN0521) and cDNA reverse transcribed with SuperscriptIII (ThermoFisher Scientific, 18080-093) while using oligo(dT)_16_-primer (ThermoFisher Scientific, 8080128). Quality of all prepared totalRNA-extractions was evaluated on a 5 mM Guanidinium Thiocyanate-agarose gel, for optimization purposes on a BioRad Experion system (RNA StdSens Assay, 7007153, 7007154) and totalRNA for sequencing was run on a 2100 Bioanalyzer Instrument (Agilent Technologies, 5067-1511). qPCR reactions were set as technical triplicates with KAPA SYBR FAST qPCR Master Mix (KR0389, v9.13) including 200 nM final concentration of forward and reverse primers and run on a Rotor-Gene Q 2plex HRM machine (9001550). See supplementary file 2 for list of all used qPCR primers.

### RNASeq library preparation and analysis

To avoid variability and thus confounding influences among the various RNASeq sample groups, we controlled rigorously for age of female parents, larval density to avoid larval crowding, age differences as well as developmental age itself and bacterial influences by using axenic culture conditions.

Poly(A)-containing mRNA molecules from totalRNA-samples were purified with oligo(dT)-magnetic beads, subsequently fragmented and cDNA synthesised with random primers using the TruSeq RNA Sample Preparation Kit v2. Adapter ligation and PCR-amplification precede cluster formation with a cBot cluster generation system. All samples were sequenced on a HiSeq 2500 Illumina Genome Sequencer as PE50. Reads were pseudoaligned with kallisto (v0.44.0) to a transcriptome index derived from all *Drosophila* transcript sequences of the dmel release r6.19. Subsequent analysis of transcript abundances was performed in R with sleuth (v0.30.0) including principal component analysis for dimensionality reduction, statistical and differential expression analysis based on the beta statistic derived from the wald test. Enriched gene ontology terms were identified by calculating hypergeometric p-values via the GOstats (v2.48.0) R package. Gene IDs were converted via the AnnotationDbi (v1.44.0) package and the reference provided with the org.Dm.eg.db (v3.7.0) database and geneset intersections visualized as UpSetR plot (v1.3.3). Please, see the accompanied R markdown deposited on GitHub for details (https://github.com/robertkrautz/sg_analysis).

### Immunohistochemistry

Proximal parts of staged larvae were inverted in PBS, unnecessary organs removed and samples fixed in 4% paraformaldehyde for 20 min. Subsequently, samples were washed three times for each 10 min in PBS or PBST (1% TritonX-100). Blocking was performed with 0.1% BSA in PBS (H2) or 5% BSA in PBST (anti-CC3, anti-pJNK and anti-Dorsal). Samples were then incubated in primary antibodies dissolved in blocking buffer for 12 h at 4°C or 1 h at room temperature (RT). Primary antibodies comprised rabbit anti-cleaved caspase 3 (1:200, Cell Signalling Technology, 9661), mouse anti-pJNK (1:250, Cell Signalling Technology, 9255), mouse anti-Dorsal (1:50, DSHB, 7A4) and mouse anti-Hemese (1:5; gift from István Andó). After washing as prior to blocking, secondary antibodies were applied together with DAPI (1:500; Sigma-Aldrich; D9542) and when necessary Phalloidin-546 (1:500, Molecular probes, A22283) in blocking buffer for 1 h at RT. As secondary antibodies goat anti-Mouse-IgG-Alexa546 (H+L; 1:500; ThermoFisher Scientific; A-11030) and goat anti-Rabbit-IgG-Alexa568 (H+L; 1:500; ThermoFisher Scientific; A-11011) were employed. Final washing as prior to blocking preceded dissection of the samples in PBS and separation of salivary glands. Tissues were mounted in Fluoromount-G (SouthernBiotech) and analysed with a Zeiss LSM780 confocal microscope. Images were extracted with Zen software (Blue edition) for further processing either in Adobe Photoshop CS5 Extended (v12.0.4 x64) or Inkscape (v0.92).

### Hemocyte- / pJNK- / CC3-quantification and size measurement

Pictures of stained glands were taken with a Zeiss Axioplan 2 microscope equipped with an ACHROPLAN 4x lens and a Hamamatsu ORCA-ER camera (C4742-95). Images were extracted with AxioVision software (v40V 4.8.2.0) and analysed with ImageJ. Cumulated area of hemocyte attachment, pJNK or CC3 fluorescence per SG or per SG part was filtered in the Red-channel and gland size determined by outlining glands, PPs or DPs with the ‘Polygon selection’-tool. The distribution of hemocytes in *Drosophila* can be approximated by a natural logarithm, which required transformation of hemocyte attachment- and SG-areas before calculating ratios (Sorrentino, 2010). Normality across all samples of a particular genotype and where necessary separated by gland part (i.e., proximal or distal) or time (i.e., 96 h and 120 h AED) was evaluated with the Shapiro-Wilk-test and by bootstrapping via the fitdistrplus R package (v1.0.11). Significant differences between experimental groups were determined for pairwise comparisons via the Student’s t-test after validating un-/equal variance via the Bartlett’s test. Data and statistical analysis was performed in R.

### Nuclei volume

Z-stacks of entire salivary glands were captured with a Zeiss LSM780 confocal microscope for DAPI signal. Obtained stacks were further processed in ImageJ via the ‘3D Object Counter’ plugin. Proximal and distal compartments were defined with the ‘Polygon selection’-tool. Transfer of the region of interest to all z-stack slices, signal thresholding, object identification and volume determination were integrated in a macro workflow. Data were plotted as average nucleus volume for all nuclei of individual glands or gland parts. Representative sections of individual z-stack slices showing the transition between proximal and distal compartments were cropped and added for illustration. Statistics were performed similar to the analysis of Hemocyte-, pJNK- and CC3- quantifications.

### In situ hybridization

Proximal parts of staged larvae were separated from the larval body in PBS, transferred to fixative (4% PFA), washed 4 times for 15 min in PBS and stored at -20°C in methanol. A cDNA for the Drs locus (LP03851) was obtained from DGRC. Probe synthesis and detailed ISH procedure is described elsewhere (Hauptmann, 2015). Stained salivary glands were mounted in Fluoromount-G (SouthernBiotech) and images were captured with a Leica MZ16 microscope combined with a Leica DFC300Fx camera.

## Acknowledgements

We would like to thank S. Höglund and the Imaging facility at Stockholm University for microscopy support, the Bloomington Drosophila Stock Center and the Vienna Drosophila Resource Center for fly stocks, the Drosophila Genomics Resource Center for the Drs cDNA, the Developmental Studies Hybridoma Bank for the anti-Dorsal antibody and D. Bohmann, S.Certel, Y. Engström, W.-J. Lee, B. Lemaitre as well as M. Suzanne for providing us with flies. We are especially grateful to R. Karlsson, I. Söll and G. Hauptmann for decisive feedback on experimental procedures and J. van den Ameele for his critical review of the manuscript. This work was supported by the Swedish Research Council (VR-2010-5988 and VR 2016-04077) and the Swedish Cancer Foundation (CAN 2010/553 and CAN 2013/546).

---

**Supplemental Figure 1.**
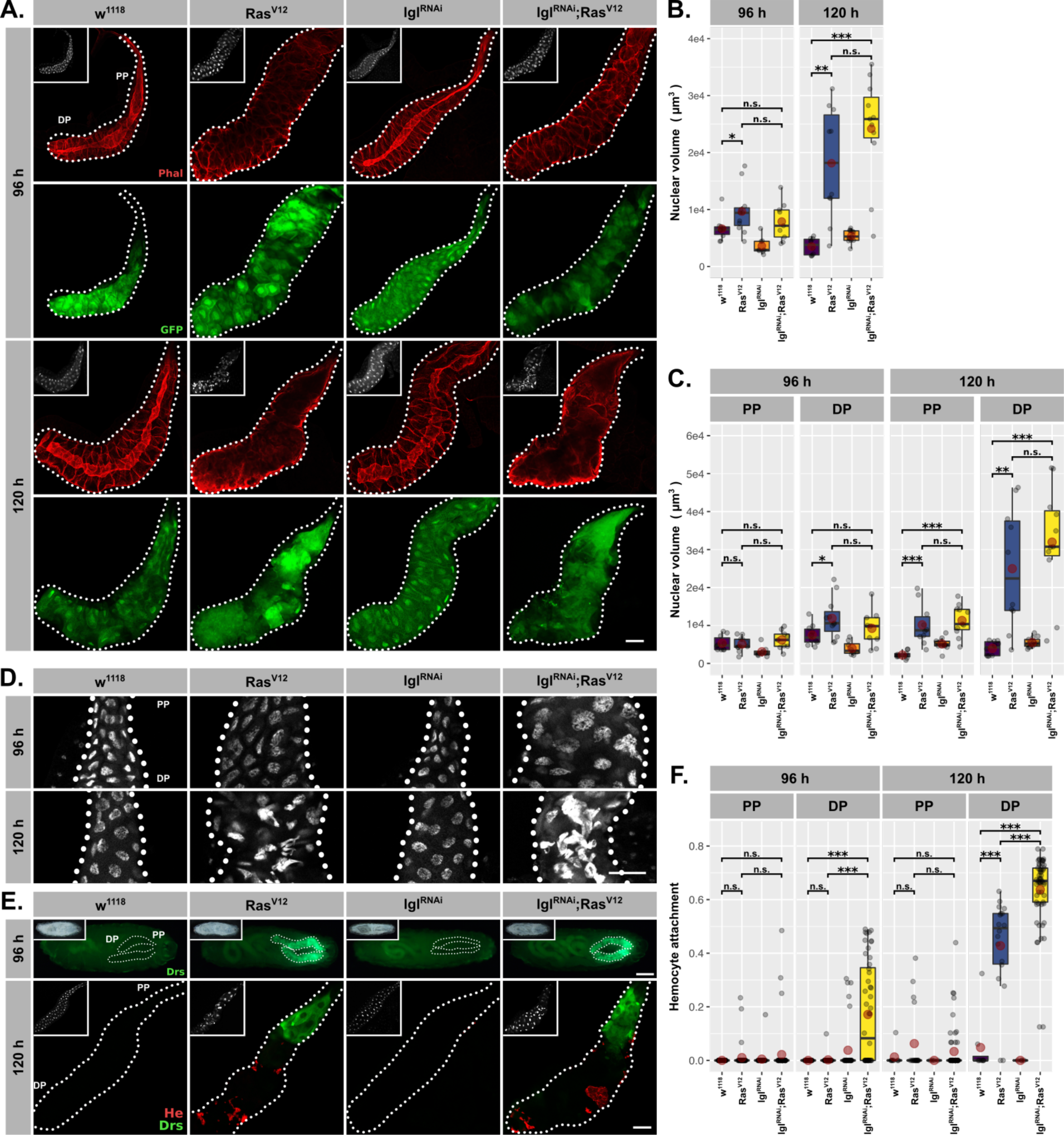
(A.) GFP (green) expression to validate *Bx^MS1096^* driver at 96 h and 120 h AED in all genotypes and Phalloidin (red) staining to trace tissue integrity. (B.) Volume of DAPI-stained nuclei derived from SG z-stacks and averaged per gland. (C.) Nuclear volume separated per gland into DP and PP. (D.) SG nuclei stained with DAPI (white) to indicate nuclear size and disintegration. (E.) Upper: Whole larvae carrying DrsGFP reporter (green). Lower: SGs with DrsGFP reporter signal (green) and stained hemocytes (anti-Hemese, red). (F.) Attached hemocytes quantified as *ln(Hemese-area)/ln(SG-area)* separated for DP and PP. Insets: (A./E. Lower) DAPI, (E. Upper) brightfield. Scalebars: (A./D./E. Lower) 100 µm, (E. Upper) 500 µm. Boxplots in (B./C./F.): lower/upper hinges indicate 1^st^/3^rd^ quartiles, whisker lengths equal 1.5*IQR, red circle and bar represent mean and median. Significance evaluated by Student’s t-tests (*** *p<0.001*, ** *p<0.01*, * *p<0.05, n.s. p≥0.05).* (A.-B./D.- E.) *w^1118^* and *Ras^V12^* data reused from Fig1. (C.) Separated nuclei measurements based on data in (B.). (F.) Separated hemocyte attachment measurements based on data in Fig1F.

**Supplemental Figure 2.**
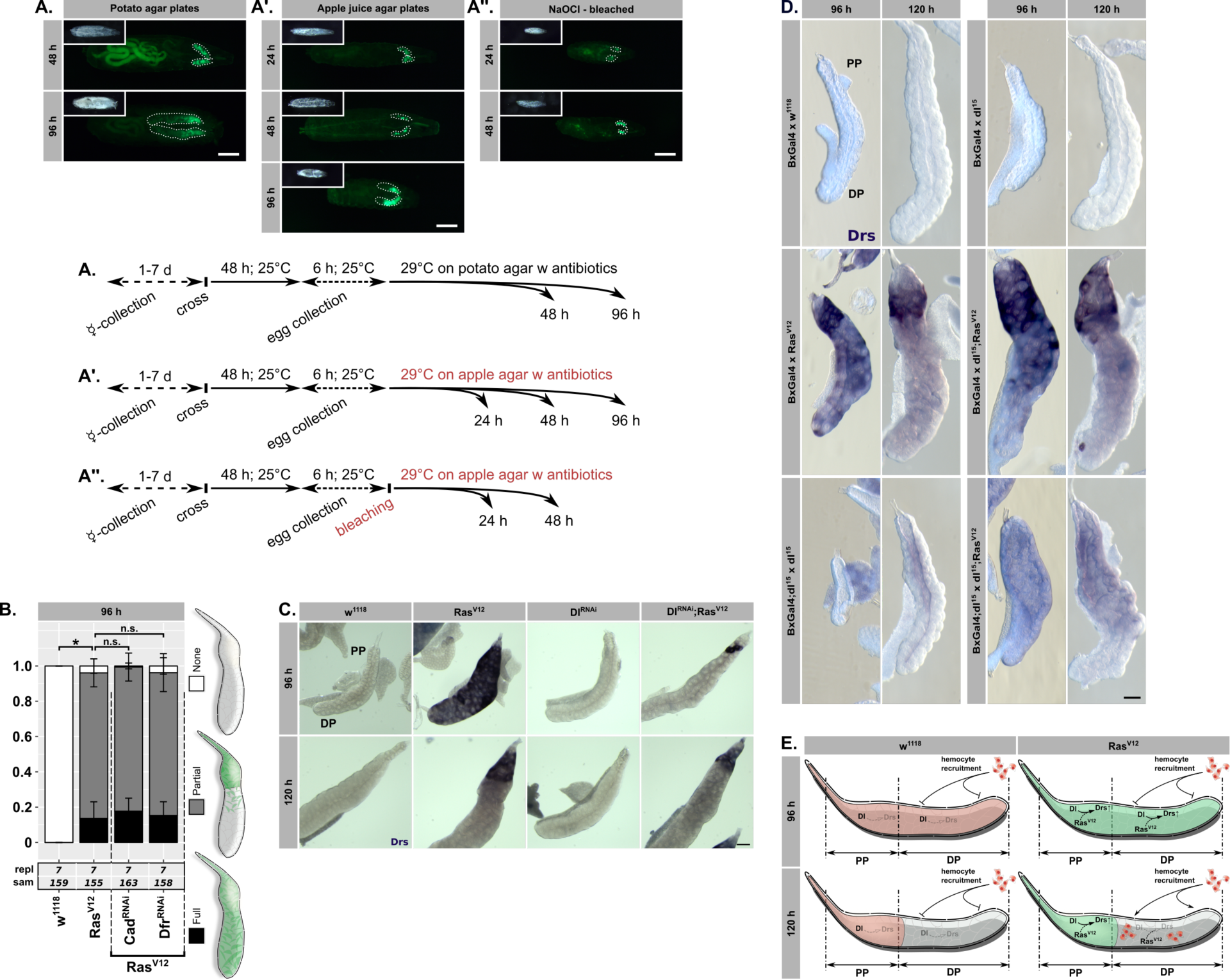
Larvae with *Ras^V12^*-glands and DrsGFP reporter (green) (A.) raised on plates with standard potatomash/molasses medium including stringent antibiotics cocktail, (A.’) transferred immediately after hatching to germ- and yeast-extract free apple-agar supplemented with antibiotics or (A.’’) after egg dechorionization were raised on sterile apple-agar including antibiotics. (B.) DrsGFP reporter assay with RNAi constructs against Caudal and Drifter in *Ras^V12^*-glands. Sketched phenotypes (right) were scored, their mean and standard deviations plotted. Dunn’s test performed to evaluate significant differences in distribution of “None”- phenotype (* *p<0.05*, *n.s. p≥0.05)*. (C.) Endogenous Drs mRNA detected by *in-situ* hybridization in *Ras^V12^*-glands with or without knocking down Dorsal. (D.) Complete set of experimental genotypes for Drs *in-situ* hybridization as shown in Fig2C including additional controls. (E.) Schematic representation of endogenous, spatio-temporally regulated Dorsal expression pattern in SGs and its interaction with *Ras^V12^* to promote Drs expression. Separation of *Ras^V12^*-glands becomes apparent at 120 h AED with Drs-expression in PP and hemocyte recruitment to DP surface.

**Supplemental Figure 3.**
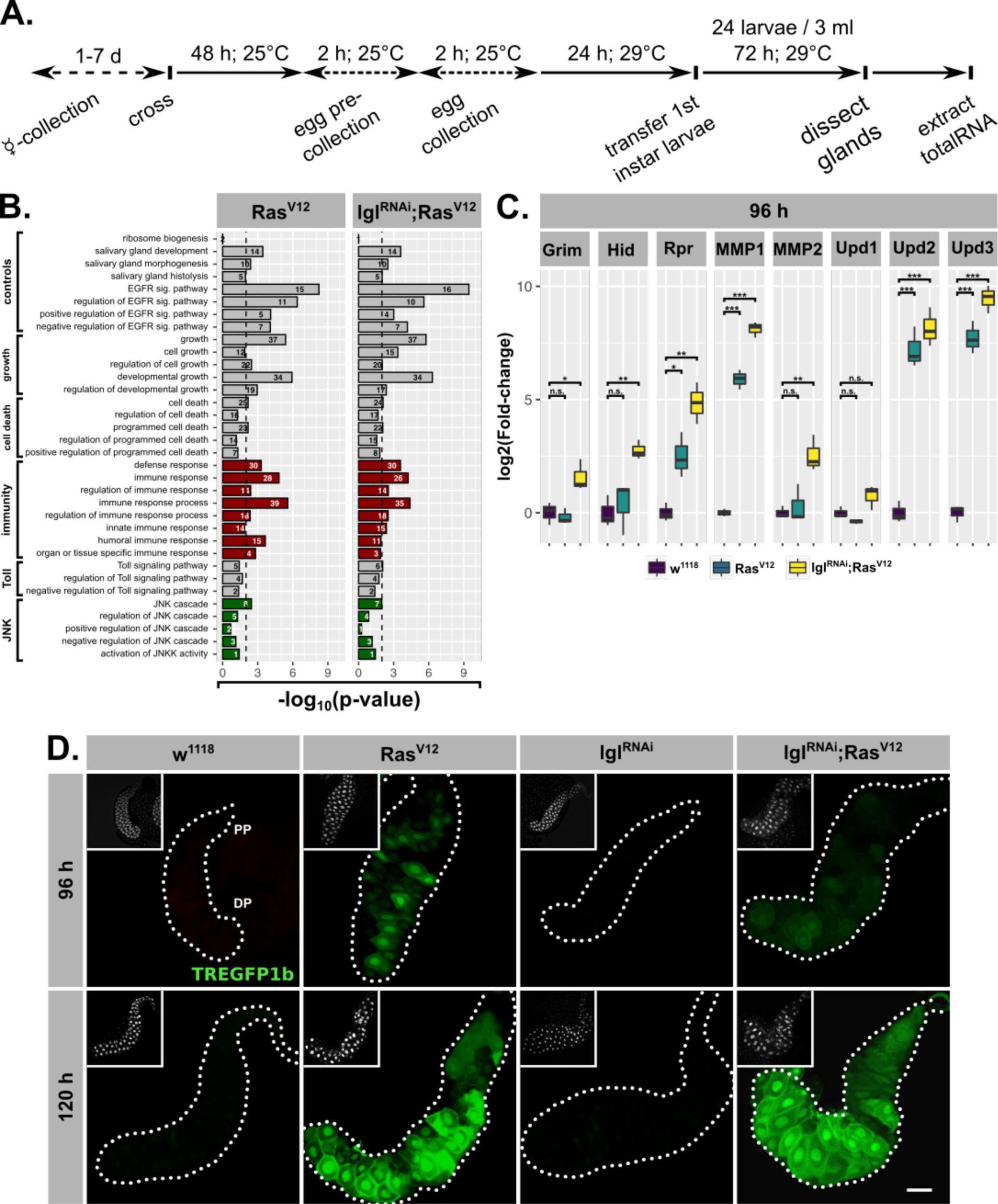
(A.) Schematic overview of protocol for preparing tissues analysed by RNAseq. (B.) Comparative GO term enrichment analysis for genes significantly upregulated in either *Ras^V12^*- or *lgl^RNAi^;Ras^V12^*-glands. Terms associated with activation of JNK (green) or immune response (red) are highlighted. GO terms for *Ras^V12^* also shown in Fig3D. (C.) Gene expression for canonical target genes measured by qPCR (*log2*-transformed, fold-change over *Rpl32*). Lower/upper hinges of boxplots indicate 1^st^/3^rd^ quartiles, whisker lengths equal 1.5*IQR and bar represents median. Significance evaluated by Student’s t-tests (*** *p<0.001*, ** *p<0.01*, * *p<0.05, n.s. p≥0.05)*. Results for MMP1, Upd2 and Upd3 in *Ras^V12^*- and *w^1118^*-glands also presented in Fig3E. (D.) Activation of TREGFP1b reporter (green) used to evaluate induction of JNK-signalling at 96 h and 120 h AED. *Ras^V12^*- and *w^1118^*-images also presented in Fig3F. Scalebar: 100 µm.

**Supplemental Figure 4.**
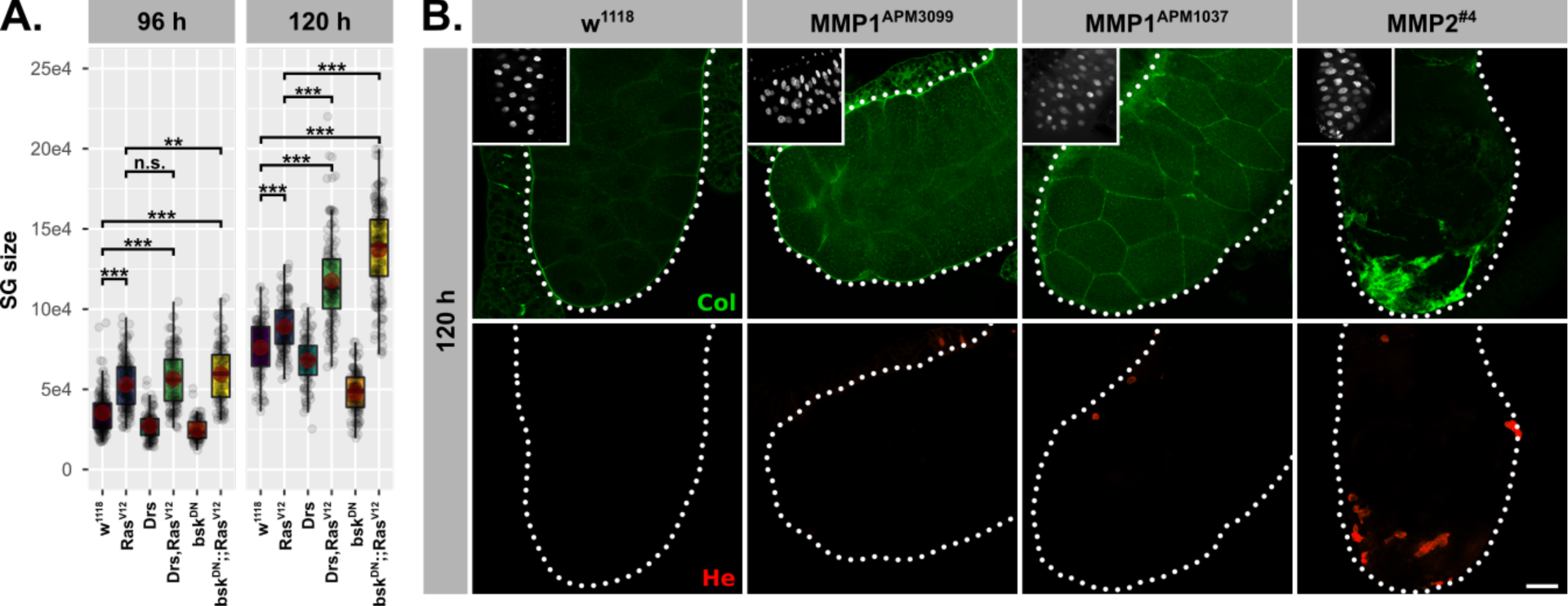
(A.) SG size measured as outlined area in captured images at 96 h and 120 h AED. Lower/upper hinges of boxplots indicate 1^st^/3^rd^ quartiles, whisker lengths equal 1.5*IQR, red circle and bar represent mean and median. Significance evaluated by Student’s t-tests (*** *p<0.001*, ** *p<0.01*, * *p<0.05, n.s. p≥0.05)*. (B.) Staining of BM via Collagen-GFP trap and attached hemocytes via Hemese antibody upon MMP1- or MMP2-overexpression. Images for MMP1^APM1037^ and MMP2^#4^ shown in Fig4E. Scalebar represents 100 µm and insets show DAPI staining.

**Supplemental Figure 5.**
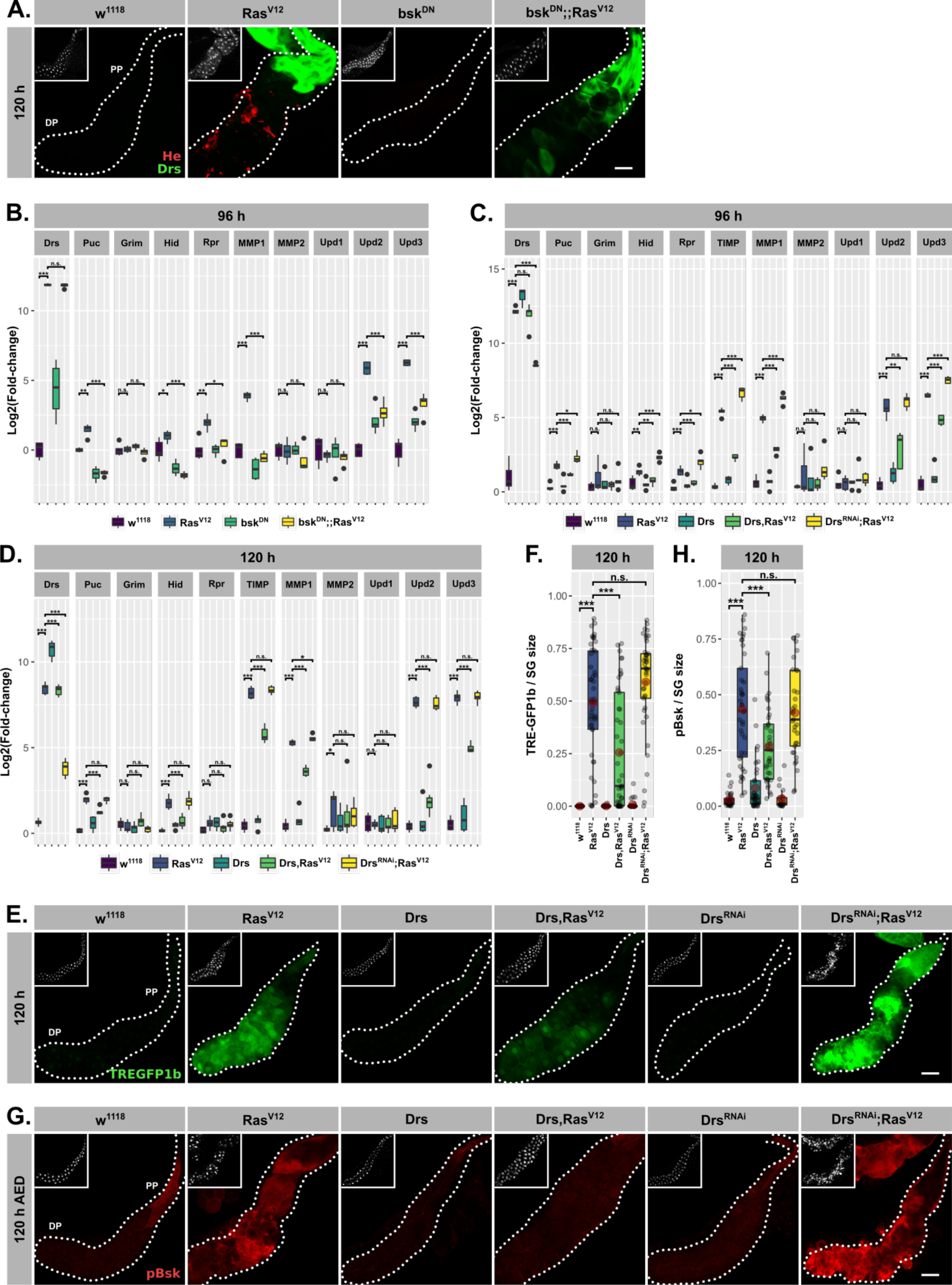
(A.) DrsGFP reporter (green) and Hemese antibody (red) used to detect endogenous Drs expression and attached hemocytes. Expression of Drs and JNK-target genes as measured by qPCR (*log2*-transformed; normalized to *Rpl32* expression) in *Ras^V12^*-glands (B.) with and without inhibited JNK-pathway at 96 h AED, (C.) with and without Drs overexpression at 96 h AED and (D.) with and without Drs overexpression at 120 h AED. *Ras^V12^*-glands with in- (Drs) or decreased (Drs^RNAi^) Drs expression (E.) including TREGFP1b reporter (green) used to detect JNK-dependent transcriptional activation and (F.) corresponding quantifications of reporter signal normalized for SG size. *Ras^V12^*-glands with in- (Drs) or decreased (Drs^RNAi^) Drs expression (G.) stained for activated basket with a phosphorylation sensitive antibody (pBsk) and (H.) corresponding quantifications of detected signal normalized for SG size. Insets: (A./E./G.) DAPI. Scalebars: (A./E./G.) 100 µm. Boxplots in (B./C./D./F./H.): lower/upper hinges indicate 1^st^/3^rd^ quartiles, whisker lengths equal 1.5*IQR, red circle and bar represent mean and median. Significance evaluated by Student’s t-tests (*** *p<0.001*, ** *p<0.01*, * *p<0.05, n.s. p≥0.05).* MMP1 and MMP2 expression in (C./D.) for *w^1118^*- and *Ras^V12^*-glands also shown in Fig4D. Hid expression in (C./D.) also presented in Fig6A. Images for *w^1118^* / *Ras^V12^* / *Drs* / *Drs,Ras^V12^* in (E./G.) also presented in Fig5E./F.

**Supplemental Figure 6.**
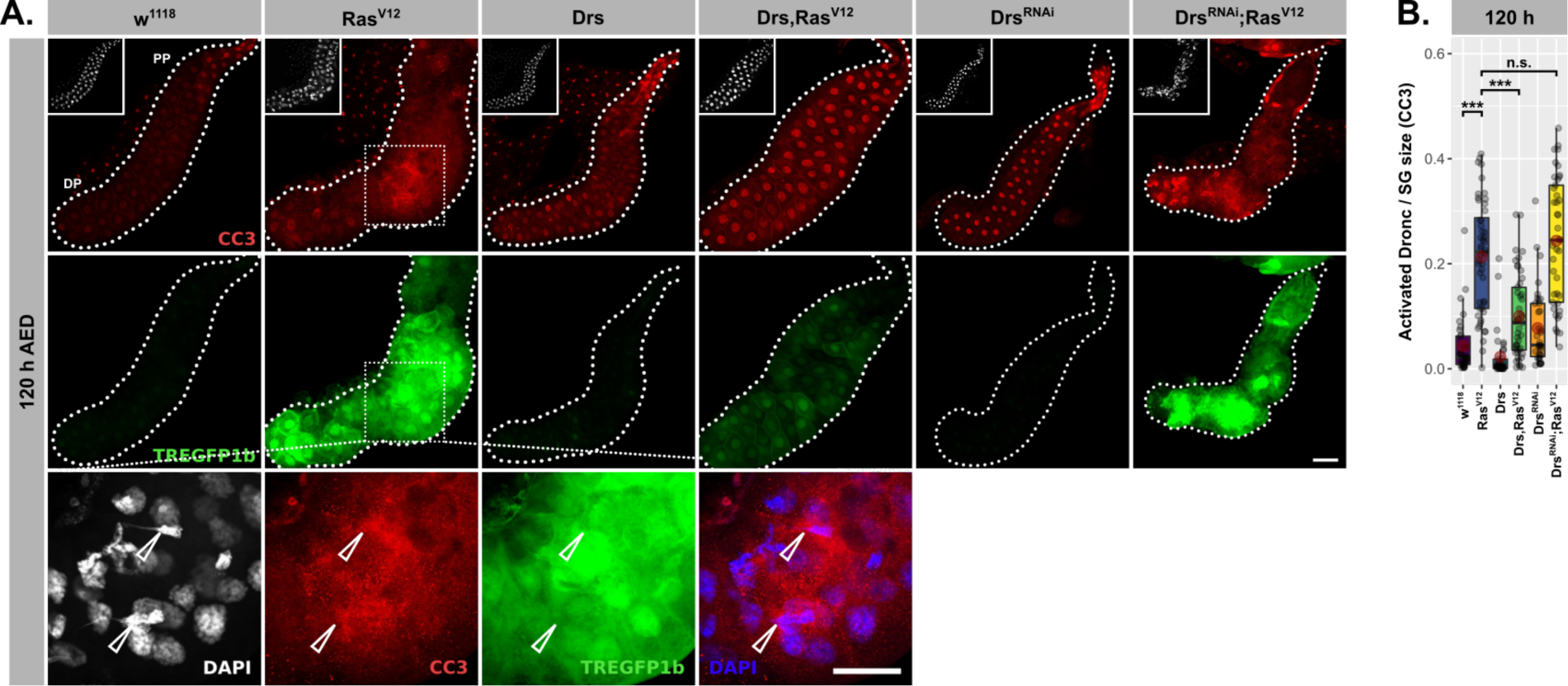
(A.) *Ras^V12^*-glands with in- (Drs) or decreased (Drs^RNAi^) Drs expression stained for activated Dronc (red) via the CC3-antibody, JNK-dependent transcriptional activation via the TREGFP1b reporter (green) and nuclei size and disintegration via DAPI (white). Arrows in magnified images point towards strong CC3-staining correlating with disintegrating nuclei. Insets represent DAPI staining and scalebars correspond to 100 µm. (B.) Quantifications of CC3-staining per gland normalized for SG size in *Ras^V12^*-glands with in- (Drs) or decreased (Drs^RNAi^) Drs expression. Lower/upper hinges of boxplots in (A./C./F.) indicate 1^st^/3^rd^ quartiles, whisker lengths equal 1.5*IQR, red circle and bar represent mean and median. Significance evaluated by Student’s t-tests (*** *p<0.001*, ** *p<0.01*, * *p<0.05, n.s. p≥0.05)*.

**Supplemental Figure 7.**
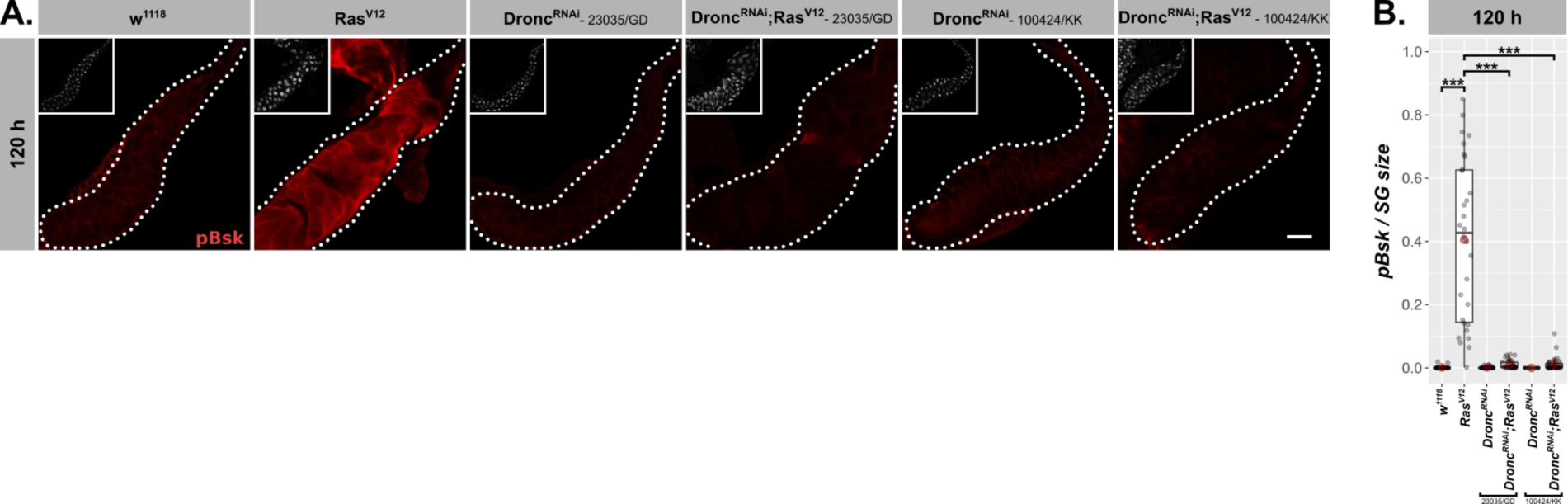
(A.) *Ras^V12^*-glands with or without Dronc-knock-down were stained for activated basket with a phosphorylation sensitive antibody at 120 h AED and (B.) the signal was quantified per SG and normalized for its size. Results for both RNAi-lines used, 23035/GD and 100424/KK, were consistent. Insets in (A.) show DAPI and scalebar represents 100 µm. Lower/upper hinges of boxplots in (B.) indicate 1^st^/3^rd^ quartiles, whisker lengths equal 1.5*IQR, red circle and bar represent mean and median. Significance evaluated by Student’s t-tests (*** *p<0.001*, ** *p<0.01*, * *p<0.05, n.s. p≥0.05)*.

